# A Living Oncogenic Microcell Isolated from Mammalian Cancers

**DOI:** 10.64898/2026.08.24.746814

**Authors:** Elena Angela Lusi, Claudia Rifici, Federico Caicci

## Abstract

The biological mechanisms underlying oncogenesis are traditionally interpreted within the frameworks of somatic mutation, clonal evolution, and, in some cases, viral infection. Here, we report the isolation, purification and characterization of a distinct class of autonomous microcellular organisms consistently recovered from independent mammalian neoplastic tissues. These entities measure approximately 1–3 μm in diameter and exhibit a reproducible organic wall assembly with internal compartmentalization revealed by transmission and scanning electron microscopy.

Biochemical and molecular analyses demonstrated a predominantly RNA-based genetic system associated with intrinsic reverse transcriptase activity. High-throughput sequencing revealed a highly distributed multipartite genetic repertoire comprising approximately 2.63 Mb organized across 1,597 independent RNA units. Canonical bacterial signatures, including 16S ribosomal RNA, were not detected, and the recovered architecture lacked the genomic organization characteristic of known retroviruses. Instead, multiple RNA units contained domains related to reverse transcriptase, mobile genetic elements, regulatory functions, and oncogene-associated sequences.

Purified preparations containing intact microcells induced rapid cellular transformation in vitro and aggressive malignancies in murine models. In contrast, matched preparations filtered through a 0.2-μm membrane failed to exhibit reverse transcriptase activity, cellular transformation, or tumorigenicity, demonstrating that the observed biological effects reside within intact micron-scale particles rather than filterable viral agents or soluble components. Furthermore, vaccination targeting the microcellular organisms was associated with tumour regression and restoration of tissue architecture in dogs with naturally occurring cancers.

Collectively, these findings describe a previously unrecognized autonomous microcell lineage within mammalian hosts possessing distinctive structural, genetic and biological properties. The combination of cellular organization, a highly distributed multipartite RNA repertoire, intrinsic reverse transcriptase activity, and oncogenic potential suggest a biological strategy not readily accommodated within current frameworks of cancer biology, virology or cellular evolution.

**Significance Statement:** We describe a previously unrecognized living oncogenic microcellular organism isolated from mammalian cancers. The organism possesses a genetic repertoire organized as numerous multipartite RNA units, exhibits reverse transcriptase activity, induces tumours in mice, and is associated with tumour regression in dogs following targeted vaccination. Distinct from both classical viruses and somatic cells, it represents an unprecedented oncogenic agent and a previously unrecognized microcell lineage not readily accommodated within current biological frameworks. The findings challenge the long-standing assumption that cancer-associated infectious agents must be small filterable viruses. More broadly, the combination of autonomous cellular organization and a distributed modular RNA genetic architecture raise the possibility that alternative strategies for storing, partitioning, and propagating biological information exist beyond the conventional genome structures recognized in contemporary biology.

**Study overview:** This investigation was conducted over approximately 10 years through collaborations with multiple independent academic laboratories, clinical centres, and contract research organizations. Key experimental findings were independently reproduced under blinded conditions. Participating institutions and technical contributions are detailed in the Acknowledgements.

**Classification:** Primary: Biological Sciences

Secondary: Medical Sciences / Microbiology/Cell Biology

## Introduction

The current taxonomy of life partitions autonomous cellular organisms exclusively into three domains: Archaea, Bacteria, and Eukarya. All known cellular lineages within these domains rely on double-stranded DNA genomes for the storage of genetic information, utilizing RNA strictly as an intermediate vector for transcription and translation. Conversely, non-cellular agents that utilize RNA as a primary genetic material—such as riboviruses and viroids—are strictly sub-micron, non-metabolic, and entirely dependent on the structural machinery of a host cell to replicate. This creates a fundamental, unchallenged dogma in modern biology: that cellular autonomy and RNA dominance are mutually exclusive.

Yet the history of biology repeatedly demonstrates that nature does not always conform to its classifications. The discovery of giant viruses blurred boundaries once thought to separate viruses from cellular organisms, while metagenomics has revealed vast reservoirs of previously unrecognized biological diversity (1–6). Such findings suggest that the architecture of life may be broader than existing taxonomic frameworks presently accommodate.

Over the course of more than a decade of investigation into human and animal cancers, we repeatedly encountered a population of micron-scale biological entities that resisted conventional classification. Initially regarded as unusual particles, these structures continued to reappear across independent specimens, different host species, and multiple experimental settings. Importantly, comparable observations were reproduced in independent research facilities operating under blinded experimental conditions. Their repeated recovery, biological stability, and reproducible isolation suggested that they represented a coherent biological system rather than a transient artifact, environmental contaminant, or degenerating host-cell structure.

The persistence of these entities across time, hosts, and experimental systems enabled a comprehensive investigation of their structural, molecular, and biological properties. The resulting observations raised questions that could not be readily addressed within existing frameworks of virology, microbiology, or cancer biology and prompted a systematic effort to determine their nature and biological significance. At the outset of this investigation, no conceptual, morphological, or molecular framework existed for the biological entities described here. This represented not simply a technical challenge, but an epistemological one. The discovery of a novel retrovirus, for example, begins within an established conceptual framework: its approximate size, morphology, replication strategy, genomic organization, and biological behaviour are already understood, allowing the newly isolated agent to be interpreted within existing biological paradigms and compared with known references.

The present investigation was fundamentally different. The biological entities described here could not initially be interpreted within any existing framework because none existed. Their morphology, developmental stages, biological behaviour, and molecular organization first had to be recognized, documented, and progressively assembled into a coherent biological concept from the experimental observations themselves. Only years later did comparisons with published descriptions of ancient cellular forms provide a broader biological context for observations that had already been established independently. The publicly available electron microscopy archive and sequencing repositories accompanying the present study now provide, for the first time, a reference framework for these microcellular organisms. Future investigators therefore begin from a fundamentally different position. Rather than constructing the biological concept de novo, they can compare newly isolated specimens directly with publicly available ultrastructural, molecular, and functional datasets, thereby substantially facilitating independent identification, validation, and reproduction of the findings reported herein.

We report the isolation, characterization, and experimental investigation of these entities using ultrastructural, molecular, biological, and functional approaches. By integrating electron microscopy, biochemical and sequencing analyses, cellular transformation assays, animal tumorigenesis studies, and therapeutic intervention experiments employing a vaccine directed against the microcellular organisms in dogs with naturally occurring cancers, we explore the possibility that they represent a previously unrecognized form of biological organization associated with mammalian cancers.

## Results

Electron Microscopy Reveals Living Microcellular Organisms Exhibiting Morphological Similarities to Early Fossil Cells

Purified preparations obtained by sucrose-gradient fractionation of human and animal cancer specimens consistently yielded discrete cell-sized entities measuring approximately 1–3 μm in diameter. Transmission and scanning electron microscopy revealed organisms bounded by a distinctive organic wall assembly (OWA) and displaying pronounced internal compartmentalization (Fig. 1). Progressive maturation stages were observed, characterized by increasing electron density and the accumulation of filamentous internal material. Reproductive activity occurred through budding, producing smaller daughter forms that subsequently matured into larger entities. The organisms were frequently detected as isolated forms or as clustered populations embedded within an electron-dense extracellular matrix.

**Figure 1.**
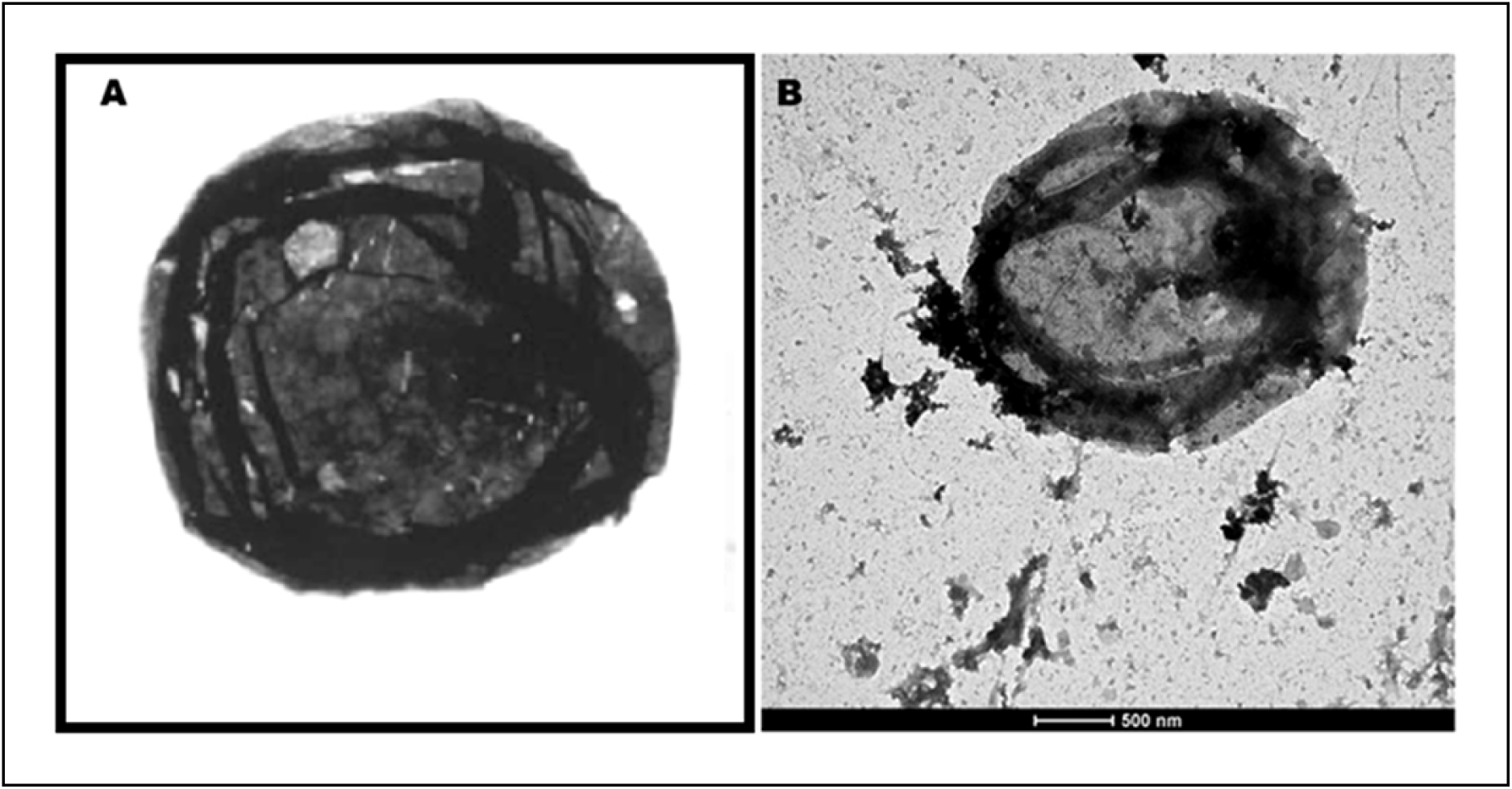
Morphological comparison between a Precambrian acritarch and a microcellular agent isolated from mammalian cancer. (A) Representative Precambrian acritarch. (B) Microcellular agent (∼3 μm in diameter) isolated from HPB-ALL human T-cell leukemia cells by sucrose-gradient purification.

A reproducible sequence of developmental morphologies was observed across independent preparations, including compartmentalized intermediate stages, mature organisms displaying complex internal organization, and budding structures associated with the emergence of daughter particles. Similar developmental stages were repeatedly identified in isolates derived from distinct human and animal cancers, indicating that the observed morphology represents an intrinsic biological property rather than a specimen-specific phenomenon.

Comparison with published descriptions and tomographic reconstructions of early fossil cells revealed a striking correspondence of architectural features. Particularly notable were similarities to acritarch-type microfossils and Doushantuo embryo-like fossils, including wall organization, internal compartmentalization, cleavage-like partitioning, staged maturation, and clustered developmental forms (Fig. 2 and Supplementary Information) (18–25, 32–40). Similar correspondences were also observed with Proterozoic fungi-like microfossils and other organic-walled unicellular forms reported from Precambrian assemblages. Viewed collectively, these recurring characteristics establish a morphological pattern closely resembling cellular designs documented throughout the Precambrian fossil record. In addition to their distinctive ultrastructure, the isolated organisms consistently retained the Gram stain, a property previously reported for Mimivirus-like particles, exhibited propidium iodide-positive staining indicative of intracellular nucleic acids, and were specifically labelled by immunogold using antibodies directed against the retroviral Gag p27 antigen (Fig. 3). These complementary structural, cytochemical, fluorescence, and immunocytochemical observations provide independent evidence that the isolated organisms possess defined biological and antigenic properties extending beyond ultrastructural morphology alone.

**Figure 2.**
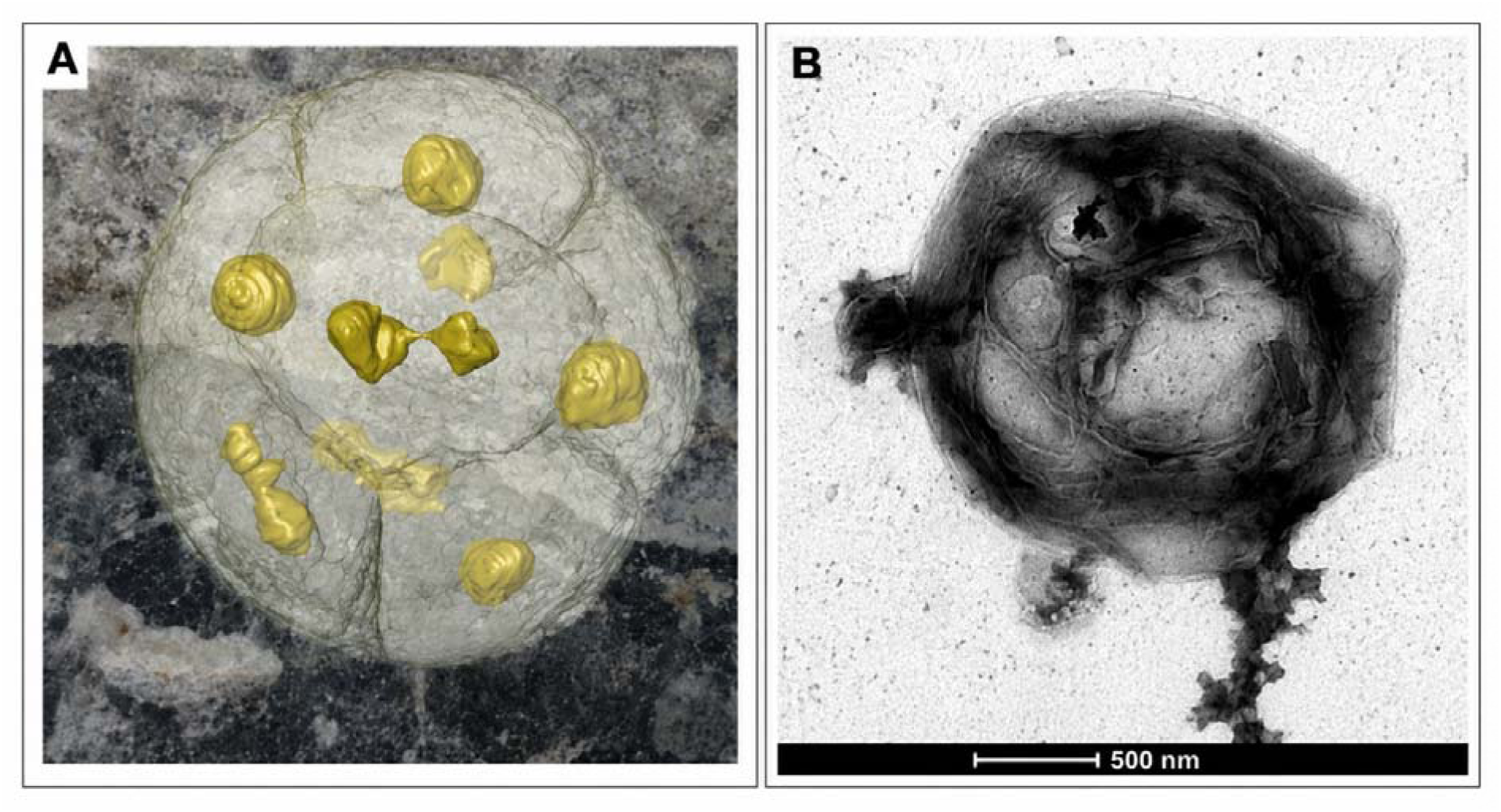
Comparison of Doushantuo fossils and the microcellular agent described in this study. (A) X-ray microtomographic reconstruction of a fossil from the Doushantuo Formation, China, showing internal partitioning and putative nucleus-like structures (yellow). Image adapted from Kaplan (2011), based on the work of Donoghue, Bengtson, and colleagues. (B) Transmission electron micrograph of a microcellular agent isolated from human cancer tissue. The organism exhibits an organic wall assembly and internal compartmentalization showing striking correspondence to the partitioned organization observed in the Doushantuo fossil. Note the similar cleavage-like subdivision pattern and central electron-dense regions.

**Figure 3.**
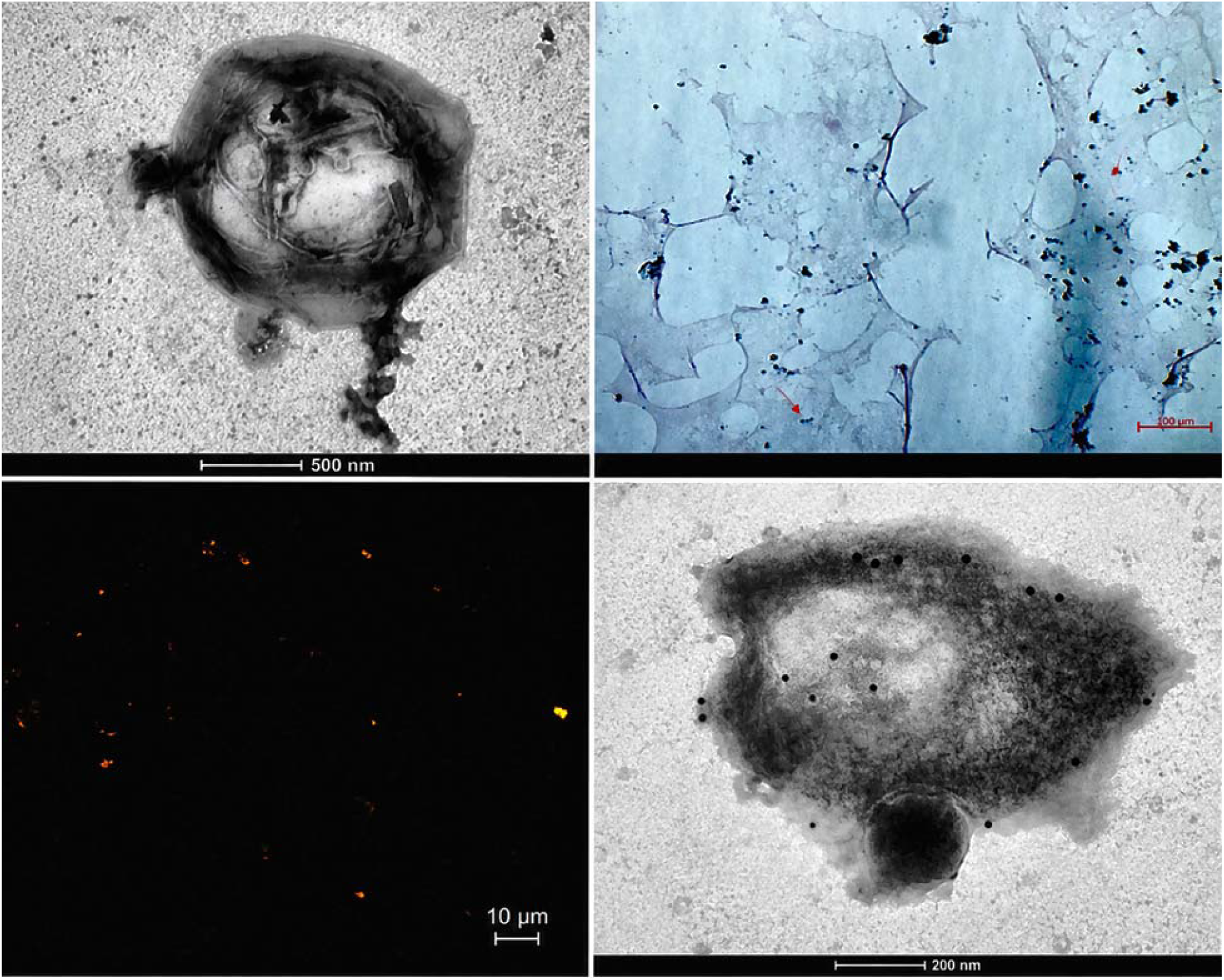
Morphological and biochemical characterization of the isolated microcellular agent. (A) Transmission electron micrograph of a representative isolated microcellular agent showing a membrane-bound structure with internal compartmentalization and complex ultrastructural organization (scale bar, 500 nm). (B) Gram-stained preparation demonstrating retention of Gram stain by the isolated microcellular particles (representative field; scale bar, 100 μm). (C) Propidium iodide fluorescence microscopy showing numerous nucleic acid-positive microcellular particles, indicating the presence of internal nucleic acid within the isolated agents (scale bar, 10 μm). (D) Immunogold transmission electron microscopy demonstrating labeling of the microcellular agent with anti-Gag p27 antibody, with gold particles localized over the particle surface and internal structures, consistent with the presence of Gag-related antigenic epitopes (scale bar, 200 nm).

To distinguish the isolated entities from common membrane-derived structures, their ultrastructural characteristics were systematically compared with those of extracellular vesicles, liposomes, apoptotic bodies, and nonspecific cellular debris. In contrast to these non-replicative or degenerative structures, the isolated microcellular organisms consistently exhibited a diameter of approximately 1–3 μm, a consistent external morphology, progressive maturation accompanied by increasing electron density, internal compartmentalization, filamentous internal material, and budding or division-like morphologies. These architectural features were reproducibly observed across independent preparations and are not characteristic of extracellular vesicles, liposomes, apoptotic bodies, or degenerating cellular fragments (Fig. 4). To facilitate independent evaluation, the complete electron microscopy archive has been deposited in a public repository listed in the Data Availability section. The dataset comprises transmission and scanning electron microscopy images acquired over more than a decade of investigation, documenting the structural diversity and developmental continuum of the isolated microcellular organisms.

**Figure 4.**
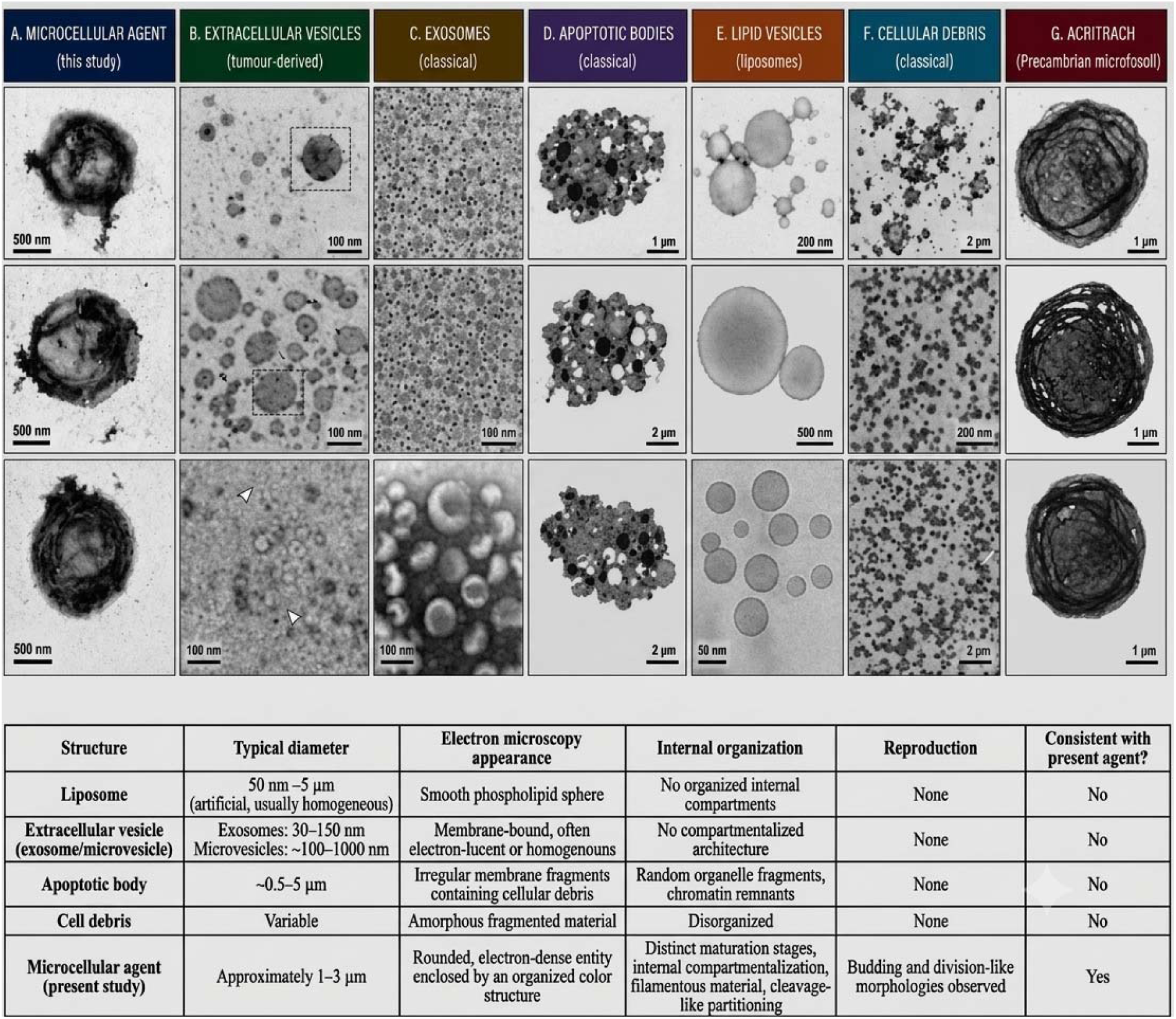
Ultrastructural comparison of the microcellular agent with extracellular vesicles, apoptotic bodies, liposomes, cellular debris, and Precambrian acritarchs. Representative transmission electron microscopy (TEM) images comparing the microcellular agent identified in this study (A, H) with tumour-derived extracellular vesicles (B), classical exosomes (C), apoptotic bodies (D), synthetic liposomes (E), cellular debris (F), and a representative Precambrian acritarch microfossil (G). The accompanying table summarizes the distinguishing ultrastructural characteristics of each entity, including size, morphology, internal organization, and reproductive features. In contrast to the comparison structures, the microcellular agent exhibits an organized outer structure, progressive maturation, internal compartmentalization, filamentous material, cleavage-like partitioning, and budding-associated morphologies. Scale bars as indicated.

Collectively, these observations identify a reproducible population of living microcellular organisms exhibiting a complex cellular architecture that closely resembles certain of the earliest cell-like forms described in the Precambrian fossil record. Occupying an unusual position between conventional viruses and somatic cells, these organisms provide the structural, cytochemical, and molecular framework for the genomic and biological characterization presented in the following sections.

## The Microcellular Organisms Exhibit Intrinsic Reverse Transcriptase Activity

Nucleic acid extraction from purified preparations revealed a marked predominance of RNA over DNA. Total nucleic acids were resistant to DNase treatment but abolished by RNase digestion, demonstrating that RNA constitutes the principal genetic material associated with the entities. DNA, when detected, was present at substantially lower abundance than RNA and may include reverse-transcribed intermediates generated by the intrinsic reverse transcriptase activity of the organisms. Reverse transcriptase (RT) activity was consistently detected in aliquots containing intact purified entities, whereas matched preparations filtered through 0.2 μm membranes lacked detectable RT activity. This segregation of activity according to particle size indicates that the observed RT activity is associated with intact microcellular organisms rather than with freely diffusible components, including filterable retroviruses, present in the surrounding medium (Fig. 5).

**Fig. 5.**
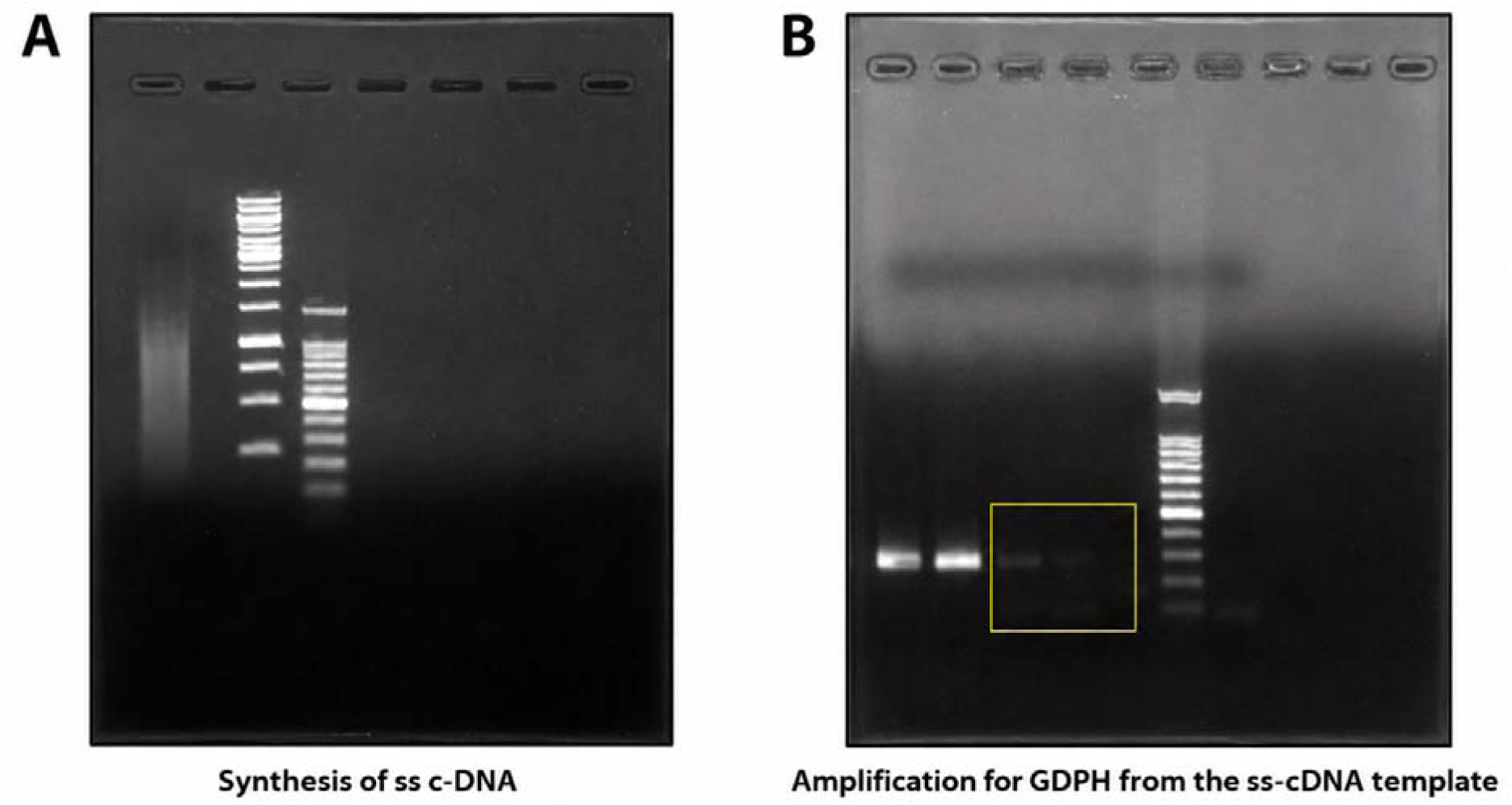
(A) Reverse transcription reaction and synthesis of single-stranded cDNA. Lane 1, reaction containing commercial reverse transcriptase; Lane 2, reaction containing lysate from purified microcellular organisms in the absence of added enzyme; Lane 3, Gene-Ruler 1-kb DNA ladder; Lane 4, 100-bp DNA ladder. (B) GAPDH amplification from cDNA templates. Lanes 1–2, reactions generated using commercial reverse transcriptase; Lanes 3–4, reactions generated using lysates from purified microcellular organisms; Lane 5, negative control; Lane 6, DNA ladder; Lane 7, additional negative control.

## RNA Sequencing Reveals a Large Multipartite RNA Genetic Repertoire Associated with the Microcellular Organisms

Sequencing of RNA-derived material obtained from purified microcellular preparations identified a complex multipartite RNA genetic repertoire comprising 1,597 independent RNA units totaling approximately 2.63 Mb of sequence information (Fig. 6). Independent sequencing runs and library preparations consistently recovered the same broad organizational features, indicating that the observed architecture represents a reproducible property of the recovered biological material rather than a dataset-specific assembly artefact.

**Figure 6.**
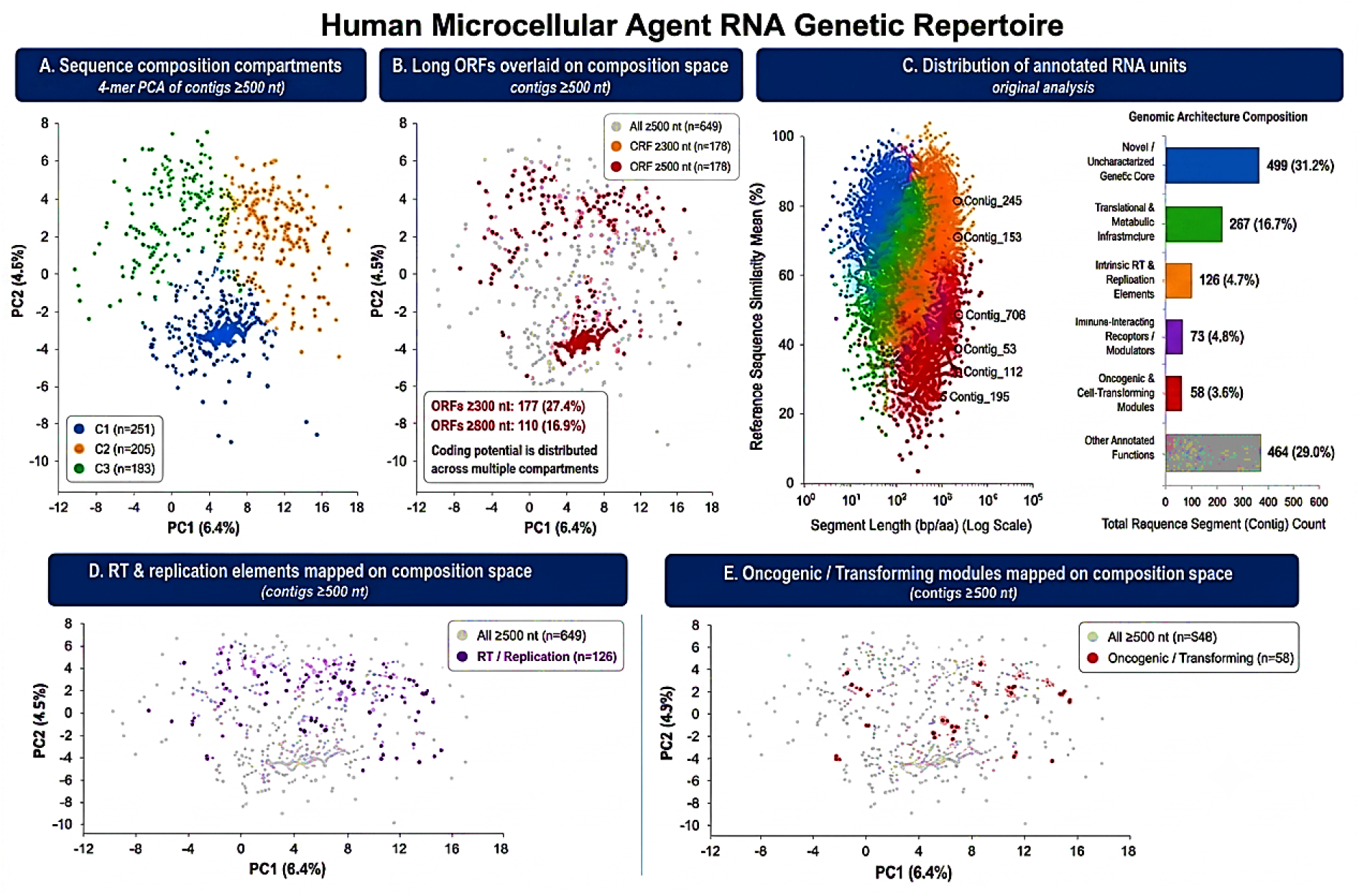
Functional landscape and compositional architecture of the human microcellular agent RNA genetic repertoire. **(A)** Principal component analysis (PCA) of RNA contigs ≥500 nt based on normalized 4-mer frequency profiles. Three major compositional compartments (C1–C3) were identified within the recovered RNA dataset. **(B)** Distribution of RNA contigs containing open reading frames (ORFs) ≥300 nt projected onto the same PCA space. Long ORF-containing contigs are enriched within specific regions of sequence space, indicating non-random distribution of coding potential. **(C)** Distribution of annotated RNA sequence elements according to sequence length and mean similarity to reference database matches. Representative sequence elements belonging to major functional classes are highlighted. **(D)** Mapping of reverse transcriptase- and replication-associated sequence elements onto the global sequence architecture. **(E)** Mapping of oncogenic and transforming sequence elements onto the same compositional framework. A total of 1,597 RNA units were recovered, of which 1,023 yielded informative annotations. The recovered sequence population exhibited structured compositional compartments, dispersed functional categories, and distributed coding potential across independent RNA units, consistent with a multipartite RNA genetic repertoire.

The RNA units ranged from 42 bp to 32.7 kb and exhibited substantial heterogeneity in sequence composition and coding potential. Rather than assembling into a single chromosomal scaffold or canonical viral genome, the sequences remained distributed across numerous independent RNA elements. This organizational pattern persisted despite repeated assembly attempts and was consistently recovered across independent analytical approaches.

Functional annotation using Blast2GO followed by targeted manual curation identified recurrent classes of biologically relevant sequences distributed throughout the assembly. These included reverse transcriptase-associated domains, integrase-like elements, kinase-associated proteins, transcriptional regulators, and sequences exhibiting similarity to oncogene families historically implicated in retroviral oncogenesis, including src-, ras-, myc-, and jun/fos-related sequence classes.

The genetic dataset did not display the characteristics expected of a conventional bacterial genome. Canonical bacterial ribosomal RNAs (16S, 23S, and 5S) were not enriched, conserved housekeeping architectures were not recovered as dominant components, and sequence composition did not converge on a profile characteristic of a single bacterial lineage. Collectively, these observations do not support interpretation of the recovered sequences as a conventional bacterial genome or a dominant bacterial contaminant.

Likewise, the dataset was not readily reconciled with a conventional mammalian transcript population. Despite the tumour origin of the starting material, dominant host-transcript signatures, coordinated tissue-specific expression patterns, and abundant repetitive host RNAs were not recovered as major components of the library. Instead, the sequence population remained dominated by orphan elements interspersed with replication-associated, regulatory, signalling, nucleic-acid-binding, and retroelement-associated functions. Although contributions from host-derived sequences cannot be excluded completely, the overall annotation profile differed substantially from that expected of simple host-cell RNA carryover.

The recovered sequences were likewise distinct from known retroviral genomes. Although reverse transcriptase-related and integrase-associated domains were repeatedly detected, no complete retroviral genomes were identified. Canonical gag-pol-env organization, long terminal repeats, and other defining features of classical retroviruses were absent. Instead, these sequence classes occurred as dispersed components distributed across multiple independent RNA units (Fig. 6).

Additional compositional analysis of the assembled sequence space failed to identify a dominant sequence family or evidence of compositional collapse. Four-mer analysis demonstrated substantial heterogeneity in sequence composition across independent RNA units, consistent with a complex distributed sequence population rather than a simple repeated contaminant. The resulting architecture was characterized by an extensive orphan component, dispersed functional categories, and the recurrent recovery of nucleic-acid-associated and retroelement-associated functions distributed throughout the dataset.

Several organizational features remained stable across independent analytical approaches. These included the persistence of extensive orphan sequence content, the inability to reconstruct a conventional contiguous genome, the absence of dominant housekeeping transcript classes, and the recurrent detection of replication-associated, regulatory, and retroelement-associated functions. Together, these recurring features indicate that the observed architecture is not dependent on a single assembly strategy or annotation pipeline but instead represents a reproducible property of the recovered sequence population.

To determine whether these organizational characteristics represented a reproducible property of the microcellular organisms rather than a dataset-specific phenomenon, an analogous analytical workflow was applied to RNA recovered from purified particles isolated from naturally occurring canine malignancies (Fig. 7).

**Figure 7.**
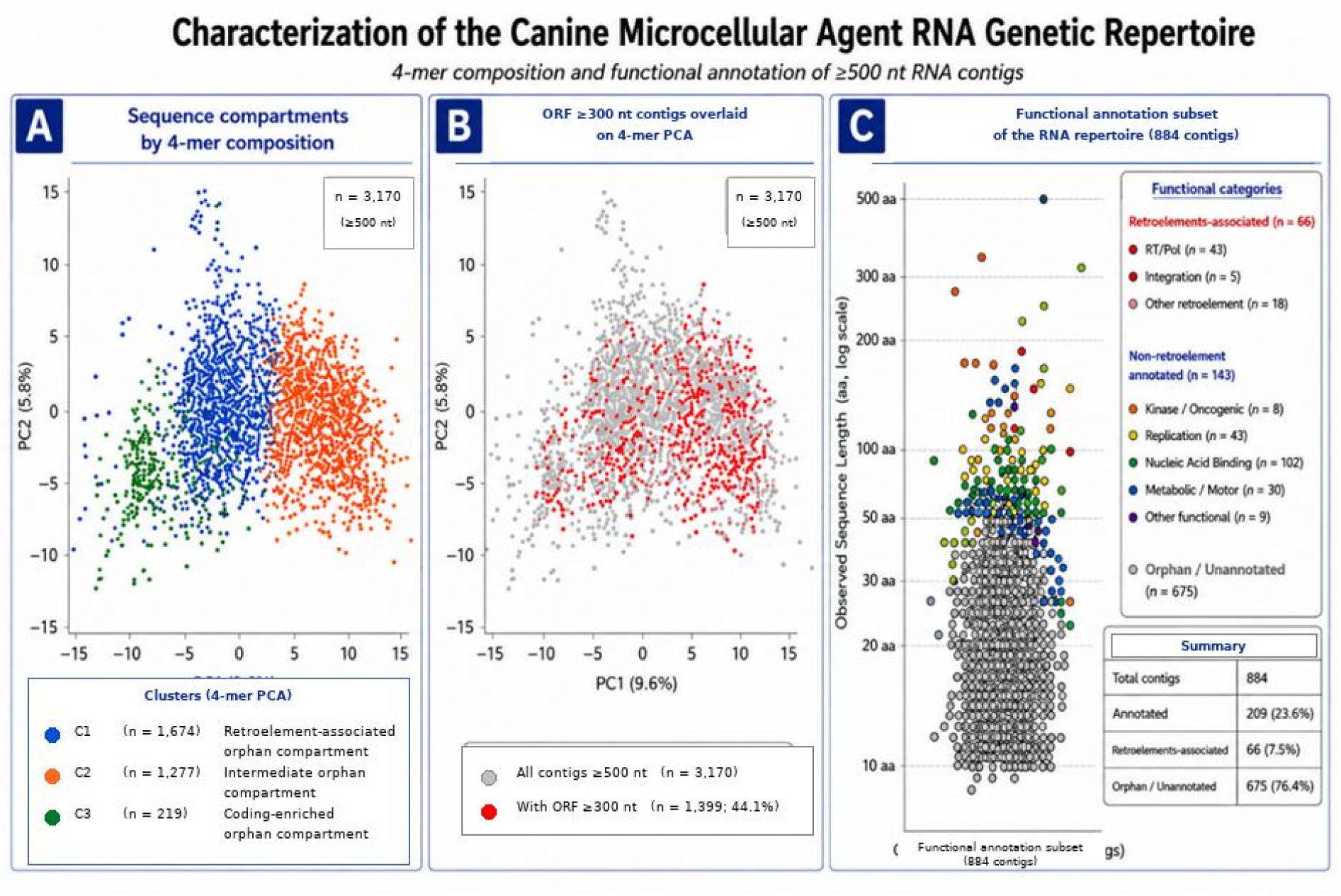
Compositional and functional architecture of the canine microcellular agent RNA genetic repertoire. (A) Principal component analysis (PCA) of RNA contigs ≥500 nt based on normalized 4-mer frequency profiles. The retained RNA dataset (n = 3,170 contigs) resolves into three major compositional compartments comprising a dominant retroelement-associated orphan compartment (C1), an intermediate orphan compartment (C2), and a smaller coding-enriched orphan compartment (C3), demonstrating a structured, non-random organization of the retained RNA sequence space. **(B)** Distribution of RNA contigs containing open reading frames (ORFs) ≥300 nt projected onto the same k-mer PCA space. Long ORF-containing contigs are preferentially concentrated within specific regions of sequence space rather than occurring randomly, indicating localized enrichment of coding potential within the structured RNA repertoire. **(C)** Functional annotation of a high-confidence subset of the canine microcellular agent RNA dataset. InterProScan-predicted functional categories (E-value ≤10⁻⁵) identified among 884 high-confidence contigs are plotted according to predicted protein length (amino acids; logarithmic scale). Replication-associated, nucleic acid-binding, metabolic, signalling, and retroelement-associated functions are distributed throughout the annotated subset, whereas orphan (unannotated) sequences constitute the predominant fraction. The annotated component therefore represents a limited functional footprint embedded within a substantially larger structured orphan RNA population. Together, these analyses indicate that the canine microcellular agent RNA genetic repertoire is organized into discrete compositional compartments exhibiting localized coding potential and a predominantly orphan sequence architecture containing a distinct retroelement-associated component.

As observed for the human microcellular organisms isolated from independent cancers, sequencing of RNA recovered from purified particles isolated from naturally occurring canine malignancies revealed a similarly distributed RNA sequence architecture that resisted reconstruction into a conventional contiguous genome. Following in silico subtraction against the Canis lupus familiaris reference genome, the resulting orphan RNA contigs were subjected to InterProScan analysis, yielding an annotated library of 884 high-confidence sequence units. This population was dominated by orphan domains (n = 675), while nucleic-acid-binding (n = 102), replication-associated (n = 43), kinase/oncogenic (n = 8), and retroelement-associated (n = 66), including integration/RNA-processing functions (n = 5), remained distributed throughout the sequence population rather than segregating into discrete functional clusters.

Notably, the canine dataset was not dominated by abundant host housekeeping, metabolic, or structural genes. Instead, replication-associated, regulatory, signalling, nucleic-acid-binding, and retroelement-associated functions remained embedded within a much larger population of orphan sequence units. The overall organization was therefore broadly analogous to that observed in the human preparations, in which diverse functional categories remained interspersed throughout a large population of independent RNA elements rather than consolidating into a conventional genomic architecture.

K-mer compositional analysis further demonstrated that the retained canine RNA library was not randomly organized. Principal component analysis of normalized 4-mer frequency profiles identified three major sequence compartments (C1–C3), while mapping of long open reading frames revealed enrichment of coding potential within discrete regions of sequence space. Functional annotation remained broadly dispersed throughout the collection, and a distinct retroelement-associated sequence footprint was embedded within a substantially larger orphan sequence population.

Taken together, these observations support the presence of a highly distributed RNA genetic complement whose organization differs fundamentally from that expected for conventional cellular genomes or classical retroviral systems. Independent analytical approaches repeatedly recovered the same broad organizational characteristics across both human and canine preparations, including extensive orphan sequence content, dispersed functional categories, structured sequence compartments, and the absence of a reconstructable contiguous genome. Although the precise genetic organization of this system continues to be refined, the recurrent recovery of these features across independent analytical approaches supports the existence of a multipartite, distributed sequence framework in which coding potential is dispersed across multiple sequence compartments rather than consolidated within a conventional genome.

The human and canine datasets independently converge on the same organizational pattern: a structured population of RNA modules containing replication-associated, regulatory, signalling, nucleic-acid-binding, retroelement-associated, and oncogenic functions distributed throughout sequence space. Collectively, these observations suggest that the biological organization of the recovered microcellular organisms is encoded not within a single contiguous genome, but within the coordinated architecture of a multipartite RNA genetic repertoire. Importantly, this interpretation does not depend on the precise size of the reconstructed dataset. Regardless of future refinements in sequencing, assembly, or annotation, the central observation remains unchanged: the recovered sequence space is extensive, highly distributed, internally structured, and does not exhibit the characteristics expected of a conventional host transcript population. The complete assembled RNA contigs, FASTA files, functional annotations, and associated analytical outputs have been deposited in a publicly accessible Figshare repository (Data Availability).

### Structural Profiling of SINE Elements Reveals a Non-Alu, tRNA-Derived and Phylogenetically Mosaic RNA Architecture of the Canine Microcellular Agent

To investigate the structural organization, evolutionary distribution, and potential host contribution of repetitive sequences within the recovered RNA repertoire, all assembled contigs were searched against the global SINEbase database. The analysis identified 2,039 SINE-associated transcript assignments representing 29 distinct SINE element classes. The repetitive landscape was overwhelmingly dominated by ancient Mammalian-wide Interspersed Repeat lineages: MIR, MIRb, MIRc, MIR3, and MIR1_Am accounted for 2,024 of 2,039 assignments (99.26%), whereas the remaining assignments comprised MamSINE1, AmnSINE1, and LFSINE_Vert-related elements.

Notably, no primate-specific Alu/7SL-derived SINE elements were detected among the 29 identified classes. Instead, the annotated repertoire consisted entirely of non-Alu SINE architectures associated with tRNA-derived structural heads. These elements contained SINEbase-defined tRNA-related architectures, including tRNA–CORE, tRNA–CORE–LINE, and 5S–tRNA–CORE–LINE arrangements, and were associated with promoter-head lineages related to tRNA-Gly, tRNA-Lys, tRNA-Cys, tRNA-Ala, tRNA-Arg, and tRNA-Thr. These structural organizations are characterized by internal RNA polymerase III promoter regions corresponding to Box A and Box B motifs linked to conserved central cores and, in several classes, LINE-recognition tails.

The taxonomic distribution of the SINEbase assignments was highly heterogeneous. Identified structural classes included elements originally described in cephalopods, insects and nematodes; teleost and jawless fish; cephalochordates; angiosperms; marsupials; camelids; xenarthrans; bats; hedgehogs; rodents; and broadly distributed mammalian lineages. Examples included OR2, CQ-5 and CELE45; DR-2, SlmI, AFC-2 and EbuSINE1; BflSINE1 and SB7; and WallSI4, Opo-1, Mar1, Mar3, Mac1, vic-1, DAS-IIb, VES, ERI-2 and ID-Spe. In contrast, only a minor fraction of the detected classes corresponded specifically to the host order Carnivora.

Thus, the recovered SINE profile was not dominated by a conventional canine-specific CanSINE landscape. Rather, it exhibited a broadly distributed, multi-taxonomic mosaic of ancient tRNA-derived SINE architectures, with complete absence of Alu elements and only limited representation of carnivore-specific classes. This repetitive signature is inconsistent with simple enrichment of ordinary canine host-repeat transcripts and supports the interpretation that the retained RNA population possesses a distinct modular structural organization.

### tRNA-Derived SINE Modules Suggest a Non-Canonical RNA Expression Strategy

The SINE architecture was considered in relation to the absence of canonical bacterial 16S, 23S, and 5S ribosomal RNA signatures from the recovered sequence repertoire. The predominance of tRNA-derived SINE heads identifies a distributed population of structured RNA modules containing conserved internal promoter architectures and tRNA-related sequence features. In known biological systems, structured tRNA-like RNAs and internal ribosome-entry elements can participate in RNA folding, stability, replication, translation-factor recruitment, and host-ribosome engagement.

The present sequence data do not by themselves establish the functional activity of these motifs. Nevertheless, their systematic occurrence within a multipartite RNA repertoire, including long ORF-bearing contigs, raises the possibility that the microcellular agent employs a non-canonical RNA-centred strategy for organizing and expressing genetic information. Such a strategy could involve host-ribosome recruitment, tRNA-like structural mimicry, catalytic RNA functions, alternative translation-initiation mechanisms, or other forms of RNA-mediated regulation. Direct demonstration of these activities will require structural prediction, ribosome-association studies, reporter assays, mutational analysis, and long-read sequencing.

Collectively, the SINE analysis reveals that the recovered RNA repertoire is not simply enriched in generic mammalian repeats. Instead, it contains a highly organized, non-Alu, predominantly tRNA-derived and phylogenetically mosaic structural component that may contribute to the replication, regulation, compartmentalization, or expression of the distributed RNA genetic system.

**Diagram 1.**
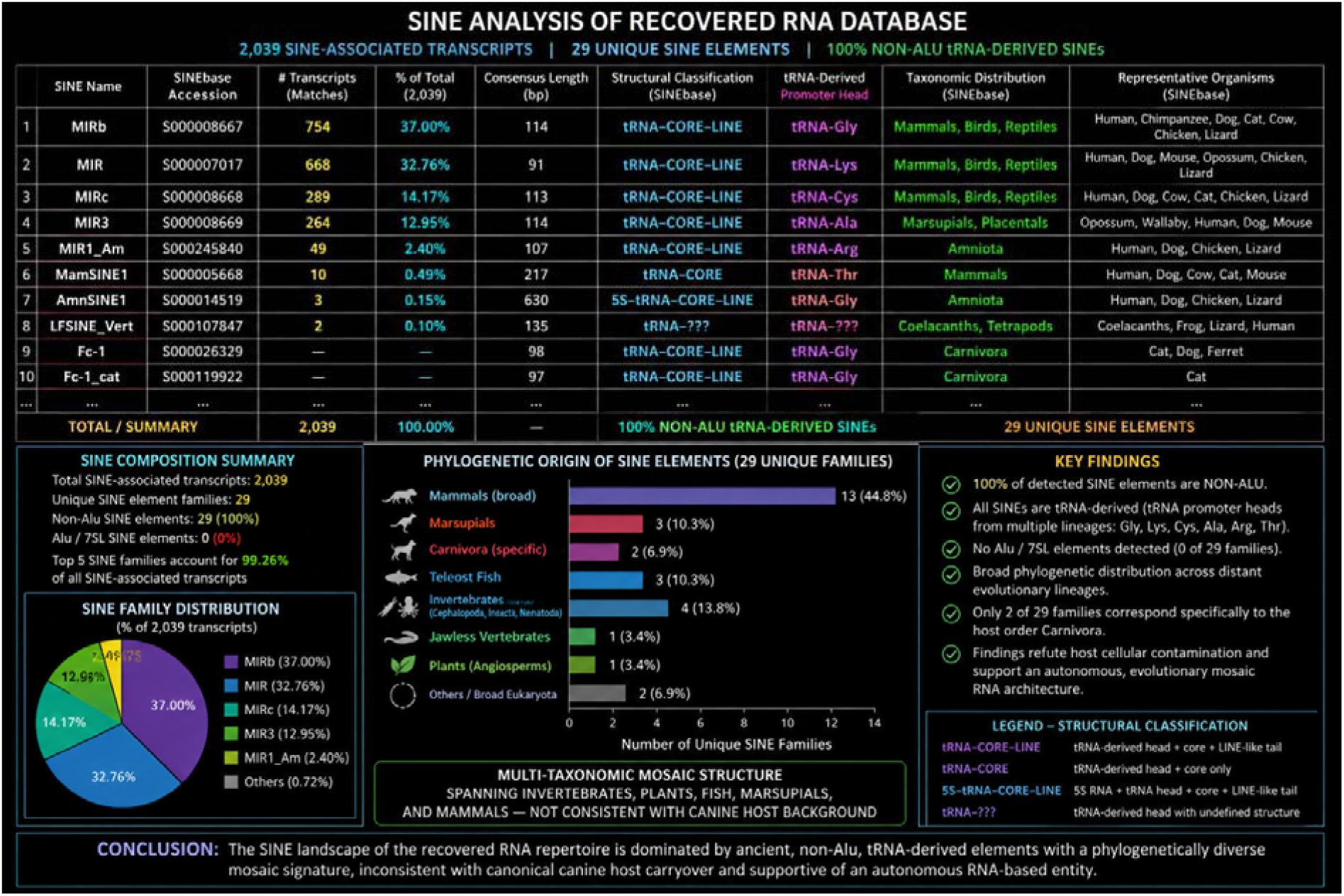
Structural profiling of SINE elements within the canine microcellular agent RNA repertoire. SINEbase analysis identified 2,039 SINE-associated transcript assignments representing 29 distinct SINE element families. The repetitive landscape was overwhelmingly dominated by ancient MIR-related elements (2,024/2,039; 99.26%), whereas no Alu/7SL-derived SINEs were detected. Structural classification demonstrated that the identified elements consisted predominantly of non-Alu, tRNA-derived SINE architectures containing conserved tRNA-associated promoter heads. Phylogenetic mapping revealed a broad mosaic distribution spanning multiple eukaryotic lineages, with only limited representation of carnivore-specific SINE families. Collectively, these findings demonstrate that the repetitive component of the recovered modular RNA repertoire is not consistent with the repetitive architecture expected from simple canine host genomic carryover but is instead enriched in ancient tRNA-derived RNA modules, revealing a structurally distinct and phylogenetically diverse repeat landscape

### Purified Microcellular Organisms Induce Cellular Transformation and Tumour Formation

Exposure of NIH-3T3 fibroblasts to purified preparations containing intact microcellular organisms resulted in rapid and reproducible cellular transformation. Within 24–48 hours, infected cultures exhibited loss of contact inhibition, increased refractility, altered morphology, and the appearance of neoplastic foci (Fig. 8). These changes progressed over time and remained stable upon serial passaging, indicating permanent transformation rather than transient cellular stress. In contrast, control cultures and cultures exposed to 0.2-μm-filtered supernatants retained normal fibroblast morphology and contact inhibition throughout the observation period.

**Figure 8.**
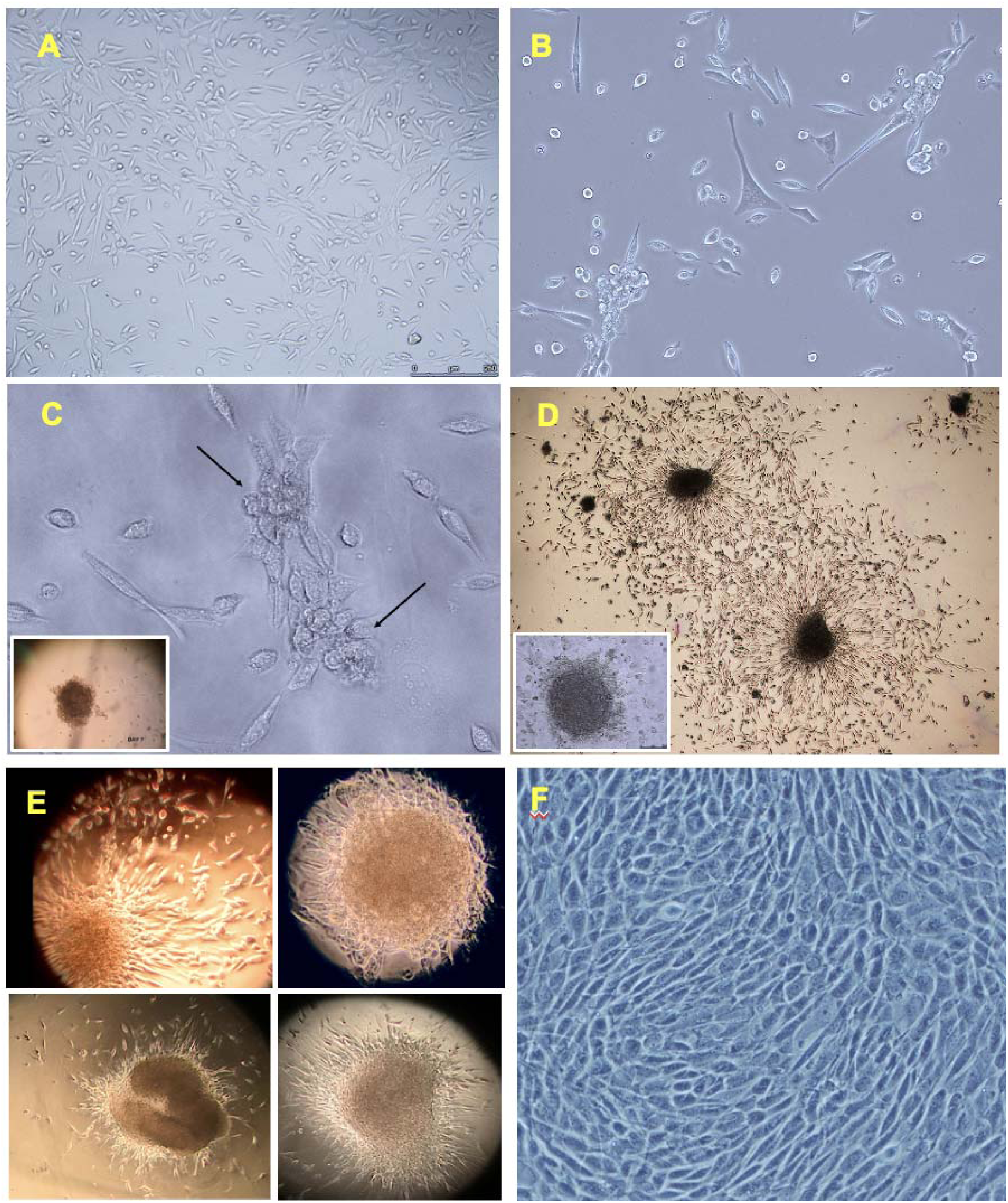
Transformation of NIH-3T3 cells by purified microcellular organisms isolated from human leukaemia cells. (A) Uninfected NIH-3T3 cells displaying normal fibroblast morphology and contact inhibition. (B) NIH-3T3 cells 48 h after exposure to purified microcellular organisms, showing altered morphology and loss of contact inhibition. (C) Early neoplastic foci (arrows) observed 48 h post-infection; inset, developing focus at day 7. (D) Progressive expansion of neoplastic foci at day 9. (E) Prominent neoplastic foci at day 10 post-infection. (F) NIH-3T3 cells exposed to matched 0.2-μm-filtered supernatant, showing normal morphology and absence of transformation. The inability of filtered supernatants to induce transformation indicates that transforming activity is associated with non-filterable cell-sized entities rather than filterable viral agents.

Transformation was characterized by the sequential emergence of discrete neoplastic foci, which expanded progressively and ultimately dominated the infected cultures. The inability of filtered supernatants to induce comparable changes demonstrates that transforming activity is associated with intact cell-sized entities and not with filterable agents present in the surrounding medium.

To evaluate tumorigenic potential in vivo, purified preparations were inoculated into immunodeficient nude mice. Infected animals developed progressively enlarging tumours at the inoculation site, followed by metastatic dissemination to distant organs. Histopathological examination confirmed the malignant nature of the resulting lesions, including highly aggressive neoplasms displaying marked cellular atypia, invasion of adjacent tissues, and frequent mitotic figures (Fig. 9 and SI). No tumours developed in animals inoculated with matched 0.2-μm-filtered supernatants.

**Figure 9.**
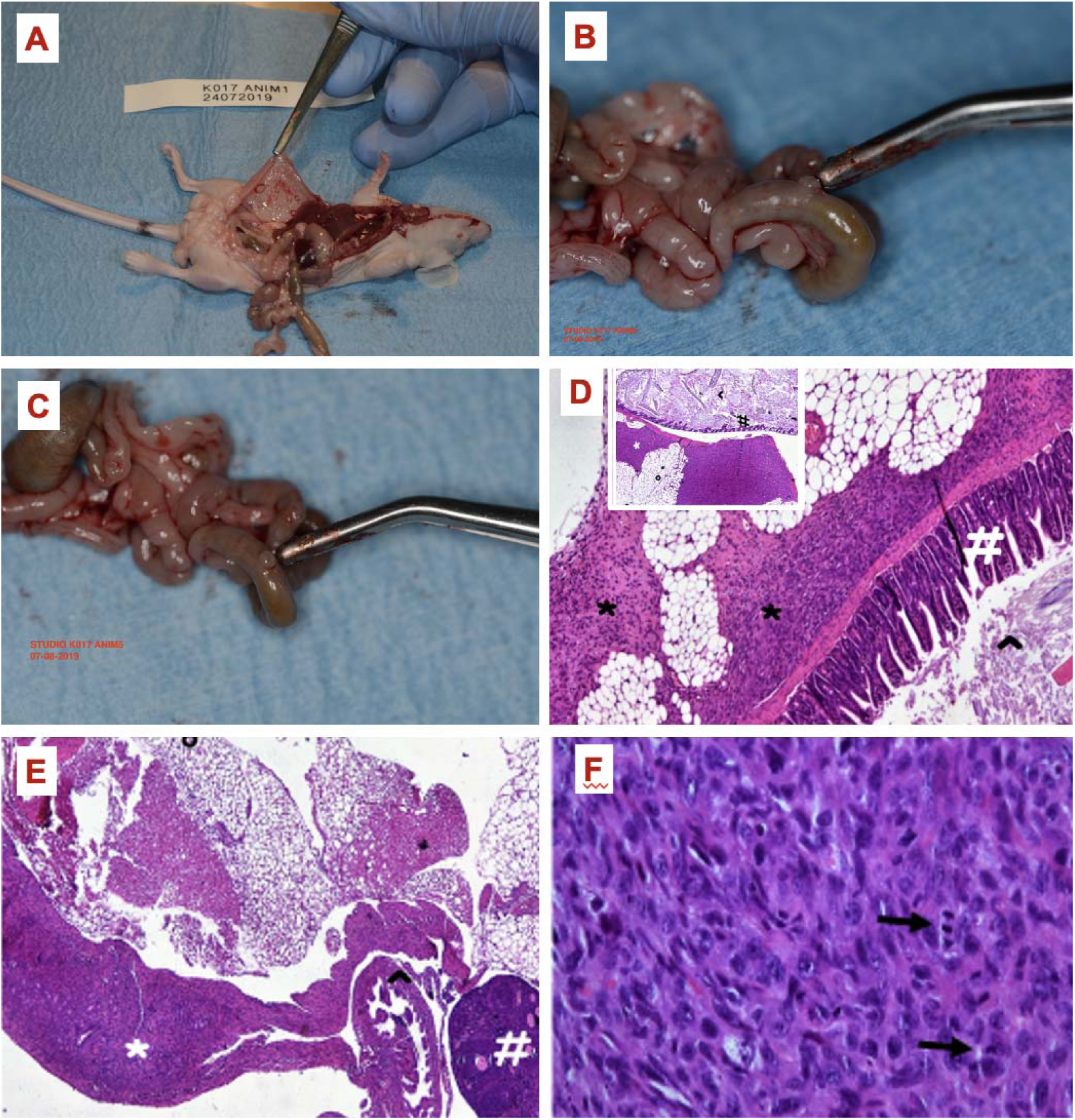
Tumorigenic and metastatic lesions induced by purified microcellular organisms isolated from human cancer cells. (A–C) Representative examples of tumours and metastatic lesions developing in mice following inoculation with purified preparations containing microcellular organisms isolated from human cancer cells. Histopathological examination confirmed the malignant nature of the resulting lesions, including peritoneal metastases and colon carcinoma observed within 4 weeks post-inoculation. (D–E) Histological sections of neoplastic lesions from infected animals. Panel (E) shows invasion of the fallopian tube by malignant tissue (*). (F) Highly aggressive anaplastic fibrosarcoma displaying marked cellular atypia and frequent mitotic figures (arrows), consistent with high proliferative activity. All lesions shown in Figure 7 developed following direct inoculation of purified microcellular organisms.

Notably, filtered supernatants derived from the same cancer specimens lacked transforming activity in vitro, failed to induce tumours in vivo, and were devoid of detectable reverse transcriptase activity. The concordant loss of transformation, tumorigenicity, and reverse transcriptase activity following filtration indicates that these biological properties co-segregate with the intact microcellular organisms. Collectively, these findings identify the purified entities as the carriers of both transforming and tumorigenic activity. The complete cellular transformation and in vivo tumourigenicity datasets, including representative images and histopathological documentation, are publicly available (Data Availability).

Vaccination Targeting the Microcellular Organisms Induces Rapid Tumour Regression and Restores Tissue Architecture

If these organisms are involved in the disease process, does experimentally targeting them produce measurable biological effects? To address this question, dogs bearing naturally occurring advanced tumours received an experimental vaccine directed against the oncogenic microcellular organisms.

Treatment was associated with rapid and reproducible tumour regression in mammary carcinomas and naturally occurring transmissible tumours, accompanied by restoration of tissue integrity, increased tumour mobility, and conversion of previously unresectable or surgically challenging lesions into clinically manageable disease (Fig. 10). Clinical responses became evident within weeks and were typically observed after two to four vaccine administrations. Ulcerated lesions progressively healed, tumours detached from surrounding tissues, and previously fixed masses became increasingly mobile.

**Figure 10.**
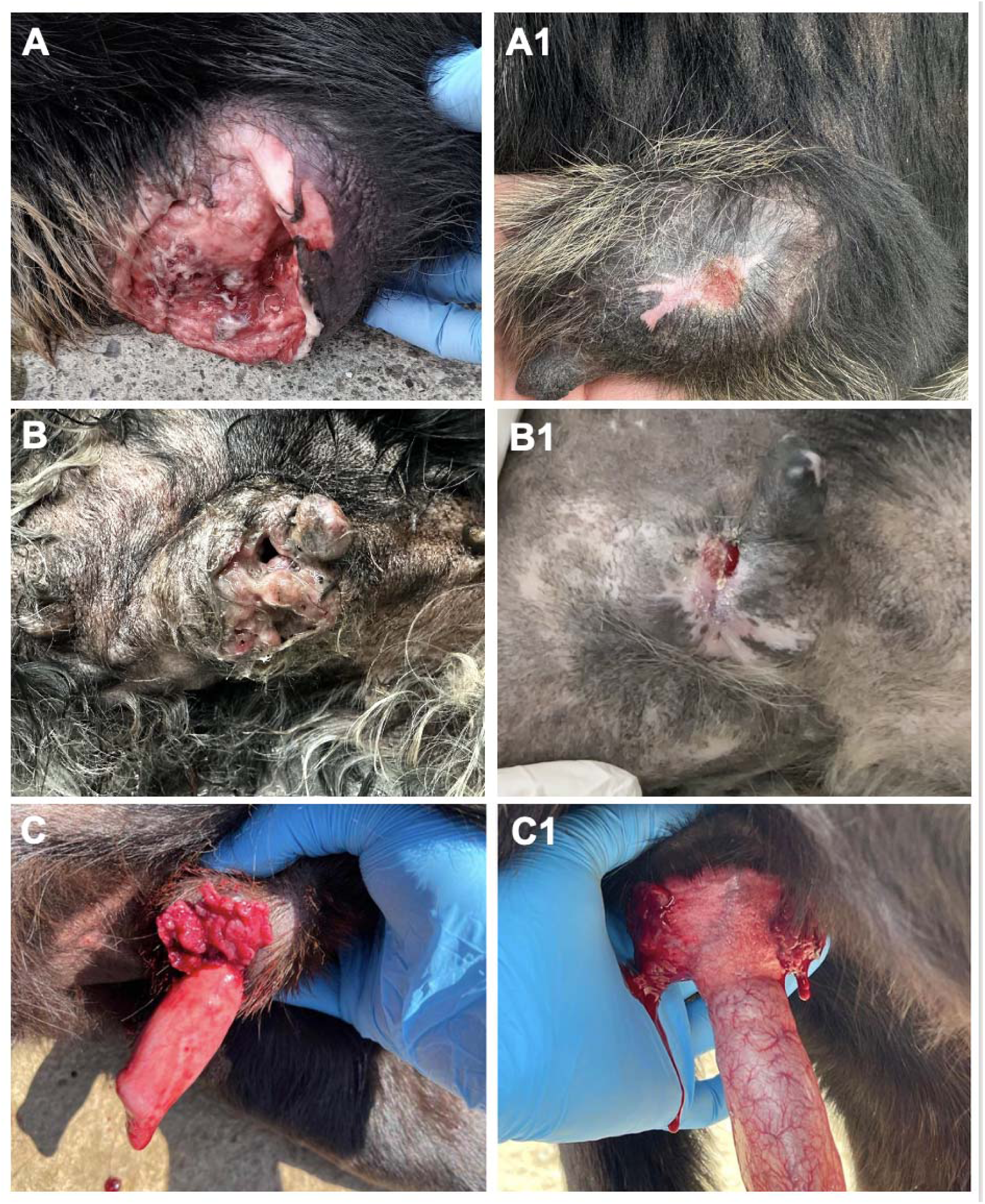
Representative examples of tumour regression and restoration of tissue architecture following vaccination directed against the isolated microcellular organisms. Case 1 (ulcerated mammary carcinoma). (A) Large ulcerated mammary tumour fixed to adjacent tissues before treatment. (A1) Marked tumour regression after four vaccine administrations, accompanied by healing of the ulcerated surface, restoration of skin integrity, and re-establishment of normal tissue planes permitting surgical excision. Case 2 (ulcerated mammary carcinoma). (B) Baseline presentation of an ulcerated, locally invasive mammary tumour. (B1) Substantial reduction in tumour volume and near-complete resolution of ulceration following vaccination. Case 3 (canine transmissible venereal tumour). (C) Naturally occurring transmissible tumour before treatment. (C1) Pronounced tumour regression and restoration of surrounding tissue architecture following vaccination.

Following repeated vaccine administration, tumour volume was substantially reduced and lesions previously considered unsuitable for surgical resection became operable owing to restoration of normal tissue planes (Fig. 10A–C1). Notably, tumour regression was accompanied by progressive re-establishment of tissue integrity and healing of ulcerated surfaces rather than fibrosis, necrosis, or destruction of adjacent tissues.

Cytological examination revealed a marked reduction in tumour cellularity together with nuclear condensation, fragmentation, cytoplasmic vacuolization, and membrane blebbing, morphological features consistent with apoptotic cell death. Similar responses were observed across independent tumours and in naturally occurring transmissible cancers.

These findings indicate that experimentally targeting the isolated microcellular organisms is associated with measurable biological effects, including rapid tumour regression, restoration of tissue architecture, and conversion of previously unresectable lesions into operable disease, thereby providing independent functional support for the biological relevance of the isolated organisms. Although formal preclinical development and controlled clinical evaluation of the vaccine fall beyond the scope of the present discovery study, these findings are presented because they demonstrate that experimental targeting of the isolated organisms is accompanied by reproducible biological responses in vivo. Additional representative cases, histopathological analyses, and treatment responses are provided in the Supporting Information (Data Availability).

## Discussion

The identification within contemporary mammalian cancers of living microcellular entities displaying striking morphological similarities to early-life fossil cells challenges established assumptions across evolutionary biology, genomics, virology, and oncology. At first glance, such a conclusion appears improbable. Yet this study did not begin with an evolutionary hypothesis; it emerged from over a decade of observations that repeatedly resisted conventional interpretation. These biological entities, initially viewed through the lens of existing paradigms, repeatedly forced us to abandon them.

Rather than relying on any individual finding, the strength of this work rests on the convergence of independent observations across different research facilities using fundamentally distinct methodologies. Purified preparations consistently yielded intact 1–3 μm entities displaying characteristic ultrastructural features. Electron microscopy revealed organic wall structures, internal compartmentalization, cleavage-like partitioning, aggregate formation, and budding propagation. Biochemical analyses demonstrated intrinsic reverse transcriptase activity associated exclusively with intact purified preparations, whereas matched filtered fractions were consistently inactive. RNA sequencing revealed a large, highly distributed genetic repertoire organized across numerous independent RNA elements. Functional studies demonstrated rapid cellular transformation in vitro, tumour formation and metastatic dissemination in vivo, and therapeutic responsiveness to targeted vaccination.

Our present interpretation was not our initial one. Early observations suggested these entities might represent an unusual class of giant viruses. Their size and intrinsic reverse transcriptase activity prompted us to provisionally designate them "Retro-Giants" (14–17), and considerable effort was devoted to reconstructing a giant viral genome within Mimiviridae and related taxa (1–6, 10). However, accumulating evidence repeatedly contradicted this hypothesis. Unlike known giant viruses, the entities displayed a distinct biological tropism, did not require amoebal hosts, consistently lacked hallmark giant-virus genes, and failed to yield a canonical contiguous viral genome. Instead, sequencing data repeatedly resolved into a highly distributed collection of independent RNA elements. The giant-virus hypothesis was not abandoned due to insufficient investigation, but because the evidence pointed elsewhere. Nevertheless, broader conceptual lessons from giant-virus research—namely that biological complexity can transcend traditional taxonomic boundaries, that infectious agents are not confined to small filterable entities, and that unexpected biological properties, including Gram stain retention, may occur outside the domain of conventional cellular organisms—remain relevant.

The ultrastructural findings proved equally difficult to interpret because no obvious modern biological counterpart existed. The dimensions, developmental patterns, and compartmentalized organization of these entities did not fit comfortably within bacterial, viral, or eukaryotic categories. Only after systematic comparison with Precambrian microfossils interpreted as early cellular forms did a coherent picture emerge. Features initially regarded as unrelated—including organic wall assembly, cleavage-like partitioning, budding propagation, and multicellular-like aggregates—closely paralleled morphologies documented in Precambrian acritarchs and Doushantuo-type fossils (18–25, 32–40). Once recognized, these similarities provided an integrative conceptual framework. While this resemblance does not establish direct ancestry, it raises the possibility that ancient cellular strategies have persisted in unexplored biological niches.

The genomic findings were no less surprising. Conventional sequencing pipelines produced what initially appeared to be technical artifacts or incomplete assemblies: assemblies resisted closure, contigs fragmented into numerous independent elements, and the architecture failed to resemble known viral, bacterial, archaeal, or eukaryotic genomes. However, as sequencing depth increased and independent analyses converged, an alternative conclusion became unavoidable. The recovered repertoire consists of approximately 1,597 independent RNA units comprising roughly 2.63 Mb of sequence information, with replication-associated, regulatory, signalling, and oncogene-related functions distributed across these elements.

Several dataset features argue against host-cell contamination, conventional retroviruses, or giant viral genomes. Replication, regulatory, signalling, and oncogenic modules are distributed among independent RNA units rather than consolidated into a contiguous genome. More than half of the annotated sequences correspond to hypothetical proteins lacking close counterparts in model organisms. Across all functional groups, sequence similarity statistics exhibit remarkable uniformity, sharing an essentially identical evolutionary-distance profile relative to databases. Were the dataset simply a mixture of host sequences, environmental contaminants, and unrelated foreign elements, these categories would separate into distinct statistical populations. They do not. Instead, replication, regulatory, signalling, and host-like cellular functions remain tightly associated within the same evolutionary framework. This shared statistical behaviour demonstrates that the recovered RNA elements are not random contaminants, but components of a single biological system characterized by a coherent, modular, and distributed genetic architecture. This system suggests an underlying pattern in which biological information is partitioned among multiple interacting RNA units, explaining why attempts to reconstruct a linear genome failed.

A comparable pattern observed in independent canine microcell preparations indicates that this distributed architecture is reproducible across species. The structural organization of the recovered RNA repertoire is further reinforced by global SINE-base profiling. Rather than exhibiting a repetitive landscape compatible with host genomic DNA carryover or random transcript degradation, the retained sequence space is overwhelmingly dominated by ancient, non-Alu tRNA-derived structural modules organized within a multi-taxonomic evolutionary mosaic. Crucially, all annotated SINE classes correspond to SINEbase-defined tRNA-derived SINE families, which are characterized by conserved RNA polymerase III internal promoter elements (Box A and Box B motifs). At the same time, the recovered repertoire completely lacked primate-specific Alu elements and displayed only minimal representation of carnivore-specific lineages (CanSINEs). This distinct structural signature supports the interpretation that these repetitive elements are not randomly retained host relics, but rather represent integral structural components of the distributed RNA repertoire—potentially serving as non-canonical scaffolds for RNA stability, tertiary folding, and host ribosomal recruitment in the absence of self-encoded 16S/23S ribosomal machinery.

This architecture becomes particularly intriguing when viewed through the lens of RNA-world theory (7–13, 26–31). Gilbert’s original hypothesis proposed that early biological systems relied on RNA simultaneously as genetic material, catalyst, and informational intermediary (7, 9, 11, 26). Subsequent work expanded this concept to envision primitive systems where information was distributed among interacting RNA molecules rather than consolidated within a single genome. Although our findings do not prove direct descent from RNA-world organisms, they provide an experimental framework for these concepts. What initially appeared to be sequencing failure may reflect a mode of genetic organization fundamentally different from modern DNA-centred genomes, wherein RNA functions not merely as an intermediary, but as a distributed repository of biological information.

The convergence of Precambrian-like developmental morphology, intrinsic reverse transcriptase activity, and a distributed RNA repertoire is noteworthy. This combination recalls concepts advanced by Temin and later framed evolutionarily by Gilbert, in which reverse transcription bridges RNA-based and DNA-based biological systems (41). While not proving these entities are living relics of an RNA world, it suggests that biological strategies proposed for early evolution may persist in modern biology.

Importantly, a distinction must be drawn between the biological entity itself and the model used to describe its genetic architecture. The physical existence of these microcellular organisms is established through direct morphological observation, biochemical activity, experimental transmissibility, tumorigenicity, and therapeutic responsiveness. Conversely, the precise organization of their genetic repertoire remains under active investigation. Nevertheless, the repeated emergence of extensive orphan sequences, dispersed functional categories, heterogeneous compositional signatures, and resistance to linear assembly indicates a genuine multipartite biological system rather than a computational artifact. Partitioning information among multiple interacting RNA elements could confer levels of modularity, adaptability, and evolutionary flexibility unattainable within a monolithic genome.

The association between these entities and oncogenesis is supported by multiple lines of evidence. Exposure of cultured mammalian cells to purified preparations induced rapid, reproducible cellular transformation, whereas matched filtered supernatants were inactive. Inoculation of purified preparations into immunodeficient mice yielded malignant tumours and metastatic lesions, while filtered fractions lacked tumorigenic activity. Crucially, reverse transcriptase activity, transforming capacity, and tumorigenicity co-segregated exclusively with intact purified entities. The simplest interpretation linking these observations is that genetic, enzymatic, and oncogenic properties reside physically within the same biological system.

These findings also intersect with the framework established by Bishop and Varmus regarding the cellular origin of viral oncogenes (42). Unlike classical retroviruses, which incorporate captured oncogenic information into compact genomes, the entities described here harbour oncogene-associated functions distributed across numerous RNA elements within a cellular-sized structure. This raises the possibility that oncogenic information can exist in a more modular and distributed state than previously recognized—a biological architecture distinct from both retroviruses and transformed host cells.

Therapeutic intervention studies provide independent functional support for the biological relevance of the isolated microcellular organisms. Vaccination directed against these entities was associated with rapid regression, restoration of tissue architecture, healing of ulcerated lesions, and conversion of previously unresectable disease into an operable state. The reproducible biological responses observed following experimental targeting are difficult to reconcile with these organisms representing passive bystanders and instead support their active participation in the neoplastic process. Although formal development and clinical evaluation of the vaccine constitute a separate avenue of investigation, these findings provide independent functional support for the biological relevance of the isolated microcellular organisms.

The significance of this concept may extend beyond human malignancies to naturally occurring transmissible cancers, such as canine transmissible venereal tumour. Given the sparsity of endogenous retroviral elements within the canine genome, the detection of retroelement-associated sequences in the host-subtracted repertoire supports their assignment to the purified agent rather than the host background. Equally compelling, the predominance of non-Alu, tRNA-derived SINE architectures arranged within a phylogenetically mosaic repertoire argues against simple canine genomic carryover and instead reveals a distinct structural organization of the retained RNA repertoire. Rather than challenging the reality of clonality, these findings suggest that additional biological layers may exist beneath expanding tumour populations. Physically and biologically distinct from somatic cancer cells, these microcellular entities may function not as clonal units themselves, but as vectors or mobilizers of oncogenic information, contributing to the emergence, maintenance, and persistence of neoplastic programmes.

From these independent lines of evidence emerged the outline of a previously unrecognized biological category. Possessing cellular dimensions, autonomous organization, developmental complexity, intrinsic reverse transcriptase activity, and a distributed RNA genetic repertoire, these entities occupy a biological space beyond current taxonomic frameworks. Their significance extends beyond biology itself, exposing an epistemological challenge. Scientific discoveries are ordinarily interpreted within established frameworks: a newly identified virus, bacterium, or archaeon is recognized because its defining characteristics conform to existing models. Here, no such framework existed. Morphology, developmental behaviour, biochemical activity, and molecular organization had to be assembled piece by piece until a coherent biological picture emerged. The principal challenge was therefore not technical, but conceptual: recognizing that a biological phenomenon existed in the uncharted space between the archetype of a virus and that of a somatic cell—a space for which biology possessed no name and no classification.

The path was neither direct nor anticipated. Only when morphology, biochemistry, genomics, and biological activity converged did the system begin to reveal itself. Distributed RNA systems, reverse transcription as an evolutionary bridge, and mobile oncogenic modules have long existed as theoretical concepts; here they appear to converge within a single biological entity. Evolution may not have abandoned its earliest experiments, but instead carried some of them quietly into modern biology. What began as an attempt to understand an unusual oncogenic particle ultimately revealed a biological system unlike any previously described—distinct from filterable viruses, bacteria, archaea, eukaryotic cells, and clonally transmitted cancer cells. If independently validated, the importance of this work will extend far beyond the identification of a novel oncogenic agent or the development of a targeted vaccine. Its lasting significance will lie in recognizing that some of life’s earliest evolutionary strategies may still persist within contemporary biology, hidden in plain sight.

The final judgment belongs to independent verification. These entities cannot be reduced solely to sequence assemblies, annotation scores, or bioinformatic classifications. Such analyses are indispensable, yet they represent only one layer of evidence for a biological reality that exists independently of computational models. The central question is therefore not whether current analytical pipelines can readily classify these entities, but whether existing biological paradigms adequately explain them. For more than a century, infectious oncogenesis has been viewed primarily through a viral paradigm. Computational interpretations will continue to evolve; biological observations remain. Dogs do not exhibit a placebo effect. When experimental targeting of the isolated microcellular organisms is consistently associated with regression of advanced, naturally occurring cancers, bioinformatic interpretation yields to observable biological reality. A sequencing plot requires specialised interpretation; a healing ulcerated lesion does not.

## Material and Methods

### Purification of Microcellular Organisms

Fresh tumour biopsies and established cancer cell lines were washed in phosphate-buffered saline (PBS), homogenized at 4°C, and lysed by vigorous vortexing in 2.5 mL PBS containing 25 μL protease inhibitor cocktail (ABMgood, Richmond, BC, Canada). Mechanical disruption was intentionally limited to gentle homogenization and vortexing. Sonication and other harsh lysis procedures were avoided to preserve the structural integrity of the microcellular organisms. The resulting suspension was incubated at 4°C for 30 min.

Crude lysates were centrifuged at 1,100 × g (3,000 rpm) for 5 min to sediment intact cells, nuclei, large membrane fragments, mitochondria, and coarse cellular debris. The resulting clarified supernatant was carefully recovered.

Importantly, no filtration step was performed before purification. The clarified supernatant was recovered directly without prior membrane filtration, loaded onto a discontinuous 35–30–25% sucrose gradient (Sigma, Milan, Italy) in 15-mL Polyclear centrifuge tubes (Seton, USA), and centrifuged at 12,300 × g (10,000 rpm) for 2 h at 4°C. Following centrifugation, the white flocculent band localized at the 25% sucrose interface was collected and concentrated by centrifugation at 12,300 × g (10,000 rpm) for 30 min at 4°C. The same purification procedure was applied to established cancer cell lines.

This workflow deliberately differs from classical protocols developed for the isolation of filterable oncogenic viruses. Rather than selecting material on the basis of membrane filtration, the clarified particulate fraction was retained throughout purification. Filtration was performed only as a biological control after purification, allowing direct comparison between intact microcellular preparations and matched 0.2-μm-filtered fractions. This strategy follows the same conceptual principle that proved essential for the discovery of giant viruses, namely that reliance on filtration-based operational definitions may exclude biologically relevant microorganisms from subsequent analysis.

Gradient centrifugation conditions were optimized to enrich micron-scale microcellular organisms while excluding soluble components and the majority of host-derived subcellular material. Purified fractions were washed extensively to remove residual sucrose and subsequently used for histochemical staining, electron microscopy, reverse transcriptase assays, nucleic acid extraction, sequencing, cellular transformation assays, tumourigenicity studies, and additional functional investigations. Aliquots of purified preparations were cryopreserved and maintained as a long-term biological collection, allowing recovery, reinoculation, molecular characterization, and comparative analyses over more than a decade of investigation.

The isolation procedure, electron microscopy, and NIH-3T3 transformation assay were performed repeatedly throughout a longitudinal study spanning more than ten years. Purifications and biological activity assays were conducted routinely, often weekly, using independent preparations derived from multiple human and animal cancer specimens as well as established cancer cell lines. The observations reported here therefore represent reproducible findings obtained across numerous independent experimental preparations rather than the outcome of a single experimental series.

The isolation strategy described here recalls the conceptual approach that proved instrumental in the discovery of giant viruses, namely that reliance on filtration-based operational definitions may exclude biologically relevant microorganisms from investigation. Rather than selecting material according to filterability, the present workflow deliberately retains the micron-scale particulate fraction throughout purification and applies membrane filtration only as a biological control following isolation.

When this investigation began more than a decade ago, no reference datasets, ultrastructural atlases, or molecular frameworks existed for these microcellular organisms. Consequently, each isolate had to be recognized and characterized de novo through the integration of morphology, biological activity, and molecular analyses. The publicly available electron microscopy archive and sequencing repositories accompanying the present study now provide a reference framework that should substantially facilitate future identification, comparative analyses, and independent replication by other laboratories. The overall isolation strategy is summarized in the schematic workflow below.

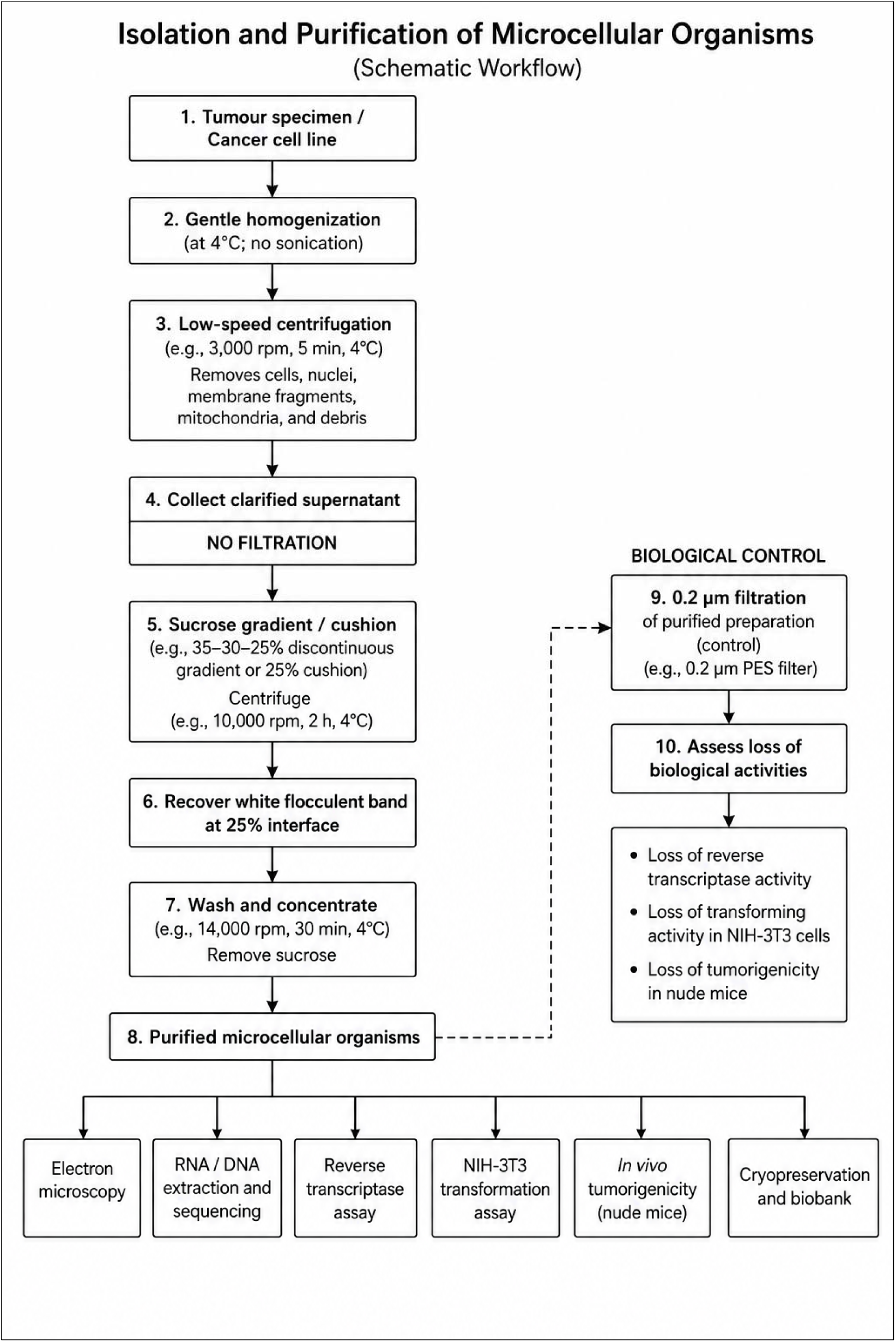

### Electron Microscopy

25 μl of purified aliquots containing the unicellular entities were placed on Holey Carbon film on Nickel 400 mesh. After staining with 1% uranyl acetate for 2 minutes the sample was observed with a Tecnai G (FEI) transmission electron microscope operating at 100 kV. Images were captured with a Veleta (Olympus Soft Imaging System) digital camera. Ultrathin sections were examined to assess size, morphology, internal organization, and aggregation patterns. Electron micrographs were obtained from multiple independent preparations to ensure reproducibility and reliability of the results. The electron microscopy image archives are available for inspection.

### RNA Extraction and Sequencing

Total RNA was extracted from sucrose-gradient-purified microcellular preparations using protocols optimized for low-input and fragmented RNA species. Residual DNA was removed by DNase treatment prior to sequencing. RNA quantity and quality were assessed using standard quality-control procedures. Sequencing libraries were prepared by the sequencing facility using Illumina-compatible library preparation methods and subjected to paired-end high-throughput sequencing. Raw sequencing reads underwent quality assessment and preprocessing prior to de novo assembly. Quality assessment was performed using FastQC. For the human dataset, de novo assembly was performed using Geneious. For the canine dataset, RNA-seq reads were assembled using the Trinity de novo transcriptome assembler. Functional annotation employed BLAST-based similarity searches, Gene Ontology mapping, and InterProScan (canine dataset), while assembly statistics were evaluated using QUAST.

### Sequence Assembly and Computational Analysis

Because no reference genome corresponding to the purified microcellular entities was available, sequencing reads were assembled de novo into contigs. Assembly of heterogeneous RNA populations can generate multiple highly similar or partially overlapping contigs representing the same underlying sequence unit. To minimize redundancy and avoid artificial inflation of repertoire size, highly similar or overlapping assemblies were clustered on the basis of sequence similarity and collapsed into non-redundant RNA sequence units for downstream analyses. The resulting non-redundant repertoire was used for annotation, compositional analyses, open reading frame prediction, and comparative organizational studies.

For the canine datasets, assembled sequences were additionally compared against the Canis lupus familiaris reference genome, and host-derived sequences were removed prior to downstream analyses. Functional annotation was performed using BLAST-based similarity searches, Gene Ontology mapping, and InterProScan domain analysis where appropriate. Particular attention was given to the identification of replication-associated, nucleic-acid-binding, signalling, retroelement-associated, and oncogenic sequence classes.

To investigate higher-order organization within the recovered sequence populations, compositional analyses based on normalized k-4 mer frequency profiles were performed. Principal component analysis (PCA) was used to identify potential sequence compartments and to evaluate whether coding potential, functional annotations, and retroelement-associated signatures were randomly distributed or concentrated within specific regions of sequence space. Open reading frames (ORFs) were predicted and mapped onto the resulting compositional landscape.

Because a substantial fraction of the recovered sequence space lacked significant similarity to currently annotated databases, both automated annotation and expert-guided manual curation were employed. Interpretation therefore incorporated not only sequence homology but also the distribution of functional classes, coding potential, compositional structure, and comparative organizational patterns independently recovered across human and canine datasets.

### Estimation of Repertoire Size

Given the fragmented and distributed organization of the recovered RNA repertoire, overall repertoire size was estimated as the cumulative length of all non-redundant RNA contigs retained after quality control and redundancy reduction. This approach is analogous to methods commonly applied to multipartite genomes, incompletely assembled giant viral genomes, and complex metagenomic systems. Repertoire size is therefore reported as cumulative sequence capacity rather than as a single contiguous genomic molecule.

For the human dataset, additional integrative analyses were performed to evaluate global organizational properties of the recovered RNA repertoire. These analyses combined sequence annotation, contig-length distributions, compositional profiling, functional-category mapping, open reading frame prediction, and distribution of replication-associated, retroelement-associated, and oncogenic sequence classes. Particular attention was given to identifying recurrent organizational patterns that remained stable across independent analytical approaches, including the presence of structured sequence compartments, extensive orphan sequence content, dispersed functional modules, and the absence of dominant bacterial or host-transcript signatures. Interpretation was therefore based not solely on individual sequence annotations but also on higher-order relationships emerging from the organization of the repertoire as a whole.

### Reverse Transcriptase Activity Assay

Reverse transcriptase (RT) activity associated with purified human microcellular preparations was evaluated using a cell-free cDNA synthesis assay. Following sucrose-gradient purification, microcellular pellets were lysed in 20 μL of lysis buffer containing 20 mM Tris-HCl (pH 7.5), 100 mM NaCl, 0.1 mM EDTA, 1 mM DTT, 50% (v/v) glycerol, and 0.25% Triton X-100 (Sigma). To assess intrinsic RT activity, 10 μL of the resulting lysate was substituted for the reverse transcriptase enzyme in a commercial cDNA synthesis kit (EasyScript cDNA Synthesis Kit, ABM) using 1 μg of Human Liver Total RNA (Thermo Fisher Scientific, Waltham, MA, USA) as template and random primers. Positive control reactions contained the commercial EasyScript reverse transcriptase enzyme, whereas experimental reactions relied exclusively on the activity associated with the purified microcellular lysates. Reverse transcription was performed at 25°C for 10 min, followed by 42°C for 50 min and enzyme inactivation at 85°C for 5 min. Following cDNA synthesis, 2 μL of the resulting cDNA was amplified by PCR using GAPDH-specific primers, Precision DNA Polymerase (ABM), 0.2 mM dNTPs, 2.0 mM MgCl₂, and 1× PCR buffer in a final reaction volume of 25 μL. PCR cycling consisted of an initial denaturation at 95°C for 5 min, followed by 40 cycles of 94°C for 1 min, 58°C for 1 min, and 72°C for 1 min, with a final extension at 72°C for 5 min. Amplification products were analysed by electrophoresis on a 1% agarose gel.

### Vaccination Protocol and in Vivo Immunological Assays

To evaluate whether purified microcellular preparations elicit targeted biological and immune responses in vivo, immunotherapeutic experiments were conducted in large animals. The principal scope and focus of this investigation remain the structural, genomic, and transforming characterization of the isolated microcellular entities; the in vivo immunological evaluations presented here serve strictly as downstream functional validations of host-entity interactions rather than a primary vaccine formulation study.

Dogs received subcutaneous administrations of an immunogenic preparation derived directly from gradient-purified microcellular organisms recovered from neoplastic tissues, formulated in a sterile, isotonic physiological vehicle buffer (pH 7.4). Individual doses were standardized to 50-100μg total protein per injection. All animals within experimental cohorts were treated according to a standardized protocol consisting of an initial priming dose followed by weekly booster injections, receiving between two and eight total doses based on clinical response kinetics. Detailed chemical compositions, specific excipient ratios, and proprietary manufacturing parameters are subject to active intellectual property protection; however, all active biological components were derived exclusively from the sucrose-gradient–purified microcellular fractions characterized throughout this work.

### NIH-3T3 Focus Formation and Transformation Assays

NIH-3T3 fibroblasts (ATCC, Lot 700009353) were maintained in Dulbecco’s Modified Eagle Medium (DMEM) supplemented with 10% fetal bovine serum (FBS). For transformation assays, 3 × 10⁵ cells were seeded into 10-cm culture dishes containing 10 mL complete medium and incubated overnight at 37°C in a humidified atmosphere containing 5% CO₂. At the time of infection, cultures were maintained at approximately 50–60% confluence. The following day, culture medium was replaced with 9 mL of fresh complete medium containing 1 mL of purified microcellular organism preparation and polybrene at a final concentration of 6 μg/mL. Cells were incubated overnight under standard culture conditions. Following a single round of exposure, the infectious medium was removed and replaced with fresh complete medium. No additional infections were performed. Uninfected NIH-3T3 cultures and NIH-3T3 cells exposed to matched 0.2 μm filtrates were used as negative controls. Cultures were monitored daily for morphological alterations, loss of contact inhibition, focus formation, and other phenotypic changes associated with cellular transformation for up to three weeks. Culture medium was replaced as required. The NIH-3T3 transformation assay was routinely employed as a biological activity assay following independent purification procedures and served as a quality-control measure throughout the study. Transformation activity consistently co-segregated with the particulate fraction containing the microcellular organisms and was reproducibly observed across multiple independent preparations over several years.

### Tumour Formation in Nude Mice

For in vivo tumorigenicity studies, six- to eight-week-old specific pathogen-free Hsd: Athymic Nude-Foxn1nu mice were obtained from Envigo RMS Srl (Udine, Italy). Animals were maintained in an isolated clean facility under controlled environmental conditions (25 ± 2°C; relative humidity 45–65%) with a 12 h light/dark cycle and provided food and water ad libitum. All experimental procedures were conducted in accordance with institutional Standard Operating Procedures and the Italian Legislative Decree No. 26/2014 governing the use of animals for scientific research. Proof-of-concept tumorigenicity experiments were performed in nude mice using two complementary approaches. In one series of experiments, NIH-3T3 cells previously transformed following exposure to purified microcellular organisms were inoculated into recipient animals. In parallel, purified preparations of the microcellular organisms were administered directly to mice to evaluate their intrinsic tumorigenic potential. As a negative control, matched 0.2 μm filtrates derived from the same preparations were evaluated and failed to induce tumour formation. Animals were monitored daily for general health status and evidence of tumour development. Gross pathological examination was performed between three-and five-weeks following inoculation. Mice were euthanized by cervical dislocation, and tissues collected at necropsy were fixed in 10% neutral-buffered formalin for histopathological analysis and downstream investigations.

### Ethics

All large animal procedures were conducted in compliance with national/international veterinary animal welfare guidelines and were approved by the Ethics Committee of the University of Messina (Protocol No. 23/2026). All procedures were conducted in accordance with institutional guidelines and applicable regulations governing clinical investigations in client-owned animals. Owner informed consent was obtained prior to enrolment for all clinical canine subjects.

## Data Availability

All data supporting the findings of this study are publicly available in dedicated Figshare repositories:

- Electron microscopy archive: https://doi.org/10.6084/m9.figshare.31802113
- Human microcellular agent RNA sequencing datasets, including assembled contigs, FASTA files, functional annotations, and associated analytical outputs: https://doi.org/10.6084/m9.figshare.32598999
- Canine microcellular agent RNA sequencing datasets, including assembled RNA contigs and compositional and functional analyses: https://doi.org/10.6084/m9.figshare.32667036
- Canine vaccination study datasets, including clinical observations, tumour-response documentation, histopathology, imaging, and supporting materials: https://doi.org/10.6084/m9.figshare.32756388
- Cellular transformation and in vivo mouse tumorigenicity datasets, including NIH-3T3 transformation experiments, tumour-formation studies, histopathology, and supporting images: https://doi.org/10.6084/m9.figshare.32620359

These repositories provide the primary experimental data and supporting analytical materials underlying the principal structural, molecular, functional, tumorigenic, and therapeutic findings reported in this study.

## Acknowledgement

This investigation originated from the observations and scientific vision of Elena Angela Lusi, who first recognized the biological entities described in this study and developed the experimental framework that enabled their subsequent isolation and characterization. Over more than a decade, she conceived, directed, and sustained the research programme, developed the experimental strategy, interpreted the findings, and provided the principal financial support that made the investigation possible.

Among all those who contributed to this work, special gratitude is owed to Federico Caicci, whose scientific integrity, generosity, and unwavering commitment made him an exceptional collaborator throughout this investigation. His expertise in electron microscopy was indispensable, and many of the ultrastructural observations forming the foundation of this study would not exist without his dedication.

Special acknowledgement is also due to Claudia Rifici, whose veterinary expertise was instrumental in coordinating the canine studies. Her generosity, courage, and intellectual openness provided invaluable support during some of the most uncertain phases of this research. She chose to stand beside this work when the data were still only whispering their story, long before the broader picture had emerged. Her confidence and commitment helped sustain a line of investigation that many would have considered too unconventional to pursue.

Deep gratitude is extended to the independent research facilities that analysed samples and generated data throughout the course of this investigation, with particular acknowledgement to VisMederi Research, Genomix4Life, and BMR Genomics. Their contributions, employing distinct methodologies and experimental approaches, were essential in establishing the reproducibility and robustness of the observations reported herein.

The bioinformatics teams who contributed to the analysis of the sequencing data are also gratefully acknowledged. In particular, we remember with appreciation Giorgio Giurato, a member of the bioinformatics group in Salerno, whose early interpretation of the sequencing data anticipated aspects of the biological framework that emerged from this work several years later. Sadly, he passed away unexpectedly before the completion of this study and before the opportunity arose to tell him how remarkably prescient many of his insights had been. His scientific insight and contribution to the bioinformatic interpretation of the sequencing data are remembered with deep gratitude.

Special thanks are due to the veterinarians, technicians, and animal-care personnel involved in the *in vivo* studies.

We also acknowledge those who challenged, questioned, and critically examined this work throughout its development. Although often difficult at the time, their scepticism compelled more rigorous experimentation, deeper analysis, and higher standards of evidence. In retrospect, these challenges strengthened the scientific foundations of the study and contributed to its eventual maturation.

Scientific discoveries are often remembered through their conclusions, but they are built through the dedication, integrity, and humanity of the people who make them possible.

## Ethics statement

All animal experiments were conducted in accordance with institutional guidelines and national regulations and were approved by the appropriate ethics committees. Experiments involving nude mice were performed using six- to eight-week-old specific pathogen-free Hsd Nude-Foxn1^nu mice obtained from Envigo RMS Srl (San Pietro al Natisone, Udine, Italy). Animals were housed under controlled environmental conditions (25 ± 2 °C; 45–65% relative humidity) with a 12 h/12 h light/dark cycle and provided food and water ad libitum. All murine procedures were performed in accordance with the Standard Operating Procedures of Emozoo, the Research Centre of Farefarma, and complied with Italian Legislative Decree No. 26/2014 governing the protection of animals used for scientific purposes. Dog vaccination experiments were approved by the Ethics Committee of the University of Messina (Protocol No. 23/2026).

## Author contributions (CRediT taxonomy)

Elena Angela Lusi: Conceptualization; Discovery; Funding acquisition; Resources; Methodology; Investigation; Data curation; Formal analysis; Ultrastructural Interpretation; Visualization; Supervision; Interpretation; Project administration; Writing – original draft; Writing – review & editing. Claudia Rifici: Investigation; Animal monitoring and follow-up; Veterinary coordination; Data curation; Validation. Federico Caicci: Investigation; Methodology; Electron microscopy; Data curation; Visualization.

